# TRaP: An Open-source, Reproducible Framework for Raman Spectral Preprocessing across Heterogeneous Systems

**DOI:** 10.64898/2026.03.26.714582

**Authors:** Yanfan Zhu, Marilyn M. Lionts, Ezekiel J. Haugen, Alec B. Walter, Trevor R. Voss, George R. Grow, Richard Liao, Meagan E. McKee, Rafay Ahmed, Joseph K. Afreh, Andrea Locke, Girish Hiremath, Anita Mahadevan-Jansen, Yuankai Huo

## Abstract

Raman spectroscopy offers a uniquely rich window into molecular structure and composition, making it a powerful tool across fields ranging from materials science to biology. However, the reproducibility of Raman data analysis remains a fundamental bottleneck. In practice, transforming raw spectra into meaningful results is far from standardized: workflows are often complex, fragmented, and implemented through highly customized, case-specific code. This challenge is compounded by the lack of unified open-source pipelines and the diversity of acquisition systems, each introducing its own file formats, calibration schemes, and correction requirements. Consequently, researchers must frequently rely on manual, ad hoc reconciliation of processing steps. To address this gap, we introduce TRaP (Toolbox for Reproducible Raman Processing), an open-source, GUI-based Python toolkit designed to bring reproducibility, transparency, and portability to Raman spectral analysis. TRaP unifies the entire preprocessing-to-analysis pipeline within a single, coherent framework that operates consistently across heterogeneous instrument platforms (e.g., Clinical Fiber-optic Raman System, Commercial Portable System and Commercial Raman Microscope). Central to its design is the concept of fully shareable, declarative workflows: users can encode complete processing pipelines into a single configuration file (e.g., JSON), enabling others to reproduce results instantly without reimplementing code or reverse-engineering undocumented steps. Beyond convenience, TRaP integrates configuration management, X-axis calibration, spectral response correction, interactive processing, and batch execution into a workflow-driven architecture that enforces deterministic, repeatable operations. Every transformation is explicitly recorded, making the full processing history transparent, inspectable, and reproducible. This eliminates ambiguity in how results are generated and ensures that identical protocols can be applied consistently across datasets and experimental contexts. Through representative use cases, we show that TRaP enables seamless, reproducible preprocessing of Raman spectra acquired from diverse platforms within a unified environment. We hope TRaP can empower Raman data processing as a reproducible, shareable, and systematized scientific practice, aligning it with modern standards for computational research. TRaP is released as an open-source software at https://github.com/hrlblab/TRaP

## 1 Introduction

Raman spectroscopy is widely used in chemistry, materials science, and biomedical research for its ability to provide molecular fingerprints without labeling or extensive sample preparation. However, raw Raman spectra typically requires multiple preprocessing steps to remove instrumental artifacts and background signals. Common processing operations include cosmic-ray removal, baseline correction, autofluorescence subtraction, smoothing, normalization, and spectral calibration.^1–4^ These procedures aim to improve spectral quality and ensure that downstream analyses reflect true chemical information rather than measurement artifacts.

The selection and ordering of preprocessing methods can influence the resulting spectral representation. Different processing strategies may produce substantially different spectral features, which can affect classification, clustering, and quantitative analysis results.^4,5^ As Raman spectroscopy becomes increasingly integrated with machine learning and large-scale spectral analysis, the need for consistent and well-documented preprocessing workflows has become increasingly important. Without standardized workflows, differences in preprocessing protocols can introduce hidden variability that complicates comparison across studies.

Another major challenge arises from variability between Raman spectroscopic systems. Differences in optical configuration, detector characteristics, excitation wavelength, and instrument response can introduce systematic variations in measured spectra. Large-scale inter-laboratory studies have shown that spectra collected using instrument configurations can vary significantly, even when measuring identical samples.^6^ To address this issue, several calibration and standardization procedures have been proposed, including Raman shift calibration standards^7,8^ and relative intensity correction protocols based on certified reference materials.^9–11^ Such procedures aim to harmonize spectral measurements across instruments and laboratories.^12–14^

Despite these algorithms and standardization guidelines, practical Raman spectral preprocessing workflows remain highly heterogeneous. In many research environments, preprocessing is implemented through custom scripts, instrument-specific software, or undocumented manual procedures, making it difficult to reproduce analyses across datasets, laboratories, or acquisition platforms. Several software tools have been proposed to support Raman spectral analysis, including graphical interfaces and open-source processing packages.^15,16^ However, these tools typically provide collections of preprocessing algorithms rather than mechanisms for enforcing standardized workflows or ensuring that processing pipelines can be reproduced across environments. At the same time, the broader scientific community has emphasized the importance of reproducible computational research and standardized data management practices. Guidelines such as the FAIR principles highlight the need for transparent and reproducible data processing workflows,^17^ while best practices stress the explicit recording of processing steps, parameters, and analysis environments.^18^ Although these principles are widely recognized, practical tools that operationalize reproducibility in Raman spectral processing remain limited.

To address these challenges, we present **TRaP** (Toolbox for Reproducible Raman Processing), an open-source framework designed to enable reproducible Raman spectral preprocessing across heterogeneous acquisition systems, as illustrated in Fig. 1. Instead of relying on ad hoc scripts or loosely connected utilities, TRaP encodes the entire processing protocol—including algorithm selection, parameter settings, and execution order—into a shareable configuration file. This configuration-driven design allows processing pipelines to be explicitly defined, stored, and reproduced across users, datasets, and computing environments. Furthermore, TRaP introduces a system abstraction layer that decouples processing workflows from instrument-specific implementations, enabling identical processing protocols to be applied consistently to spectra acquired from heterogeneous Raman platforms. The TRaP software implementation is publicly available as an open-source project at https://github.com/hrlblab/TRaP.

**Fig 1:**
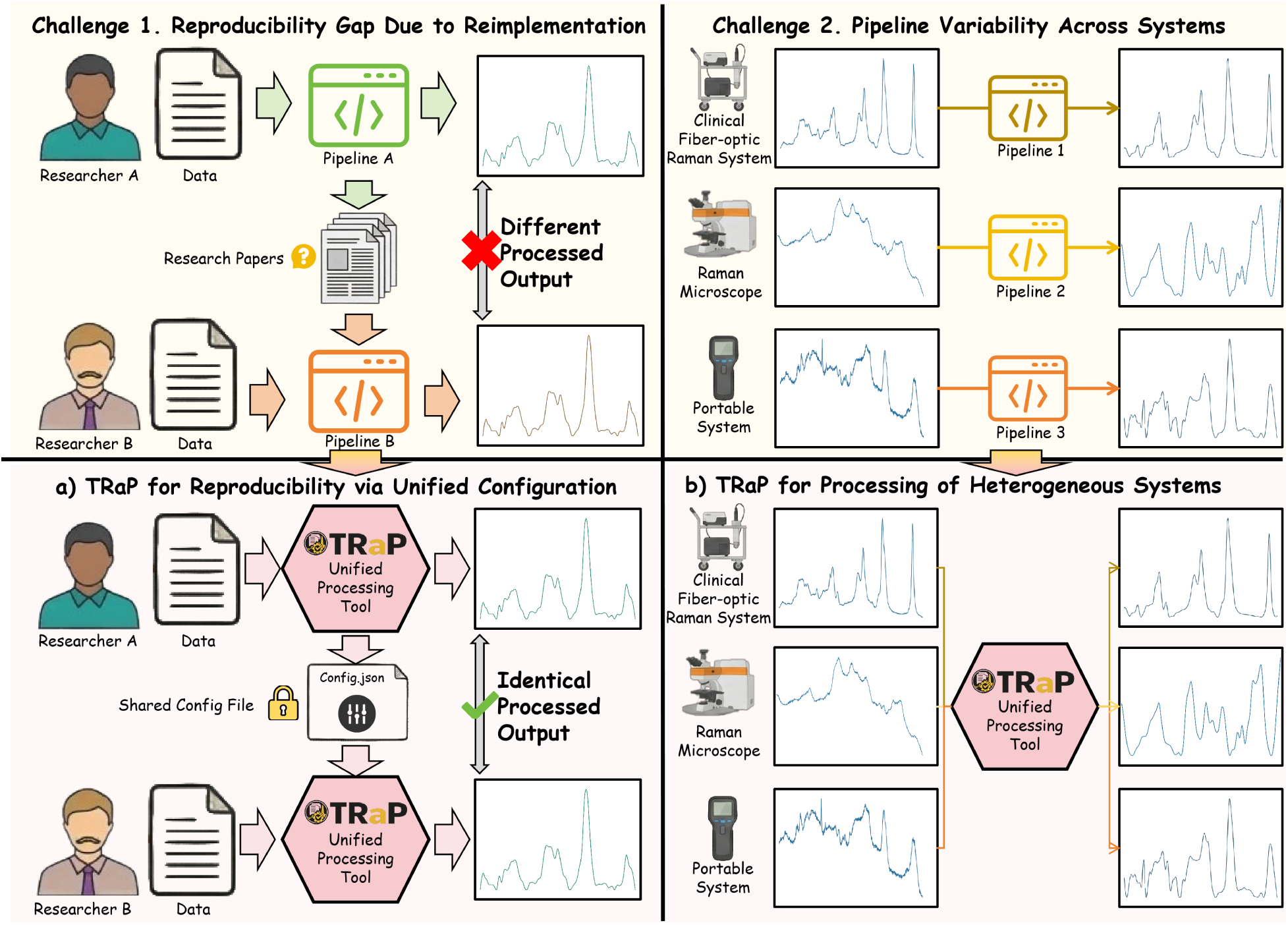
**Top-left:** Reimplementation variability—independent pipelines applied by different researchers produce inconsistent outputs. **Top-right:** System variability—heterogeneous instruments (clinical fiber-based Raman systems, commercial microscopes, and commercial portable systems) and system-specific pipelines further amplify inconsistencies. **Bottom-left (a):** TRaP ensures reproducibility via a shared configuration, enabling identical processing across users. **Bottom-right (b):** TRaP standardizes preprocessing across systems, enabling consistent and comparable outputs.

The key contributions of TRaP are summarized as follows:

- **Reproducible Raman preprocessing through configuration-driven execution.** TRaP enables exact reproduction of Raman preprocessing workflows by encoding the full processing pipeline—including algorithm selection, parameterization, and execution order—into shareable configuration files. This eliminates the need for ad hoc scripting or reimplemeting, and ensures consistent results across users, datasets, and computational environments.
- **Transparent processing with explicit provenance tracking.** TRaP provides full visibility into the preprocessing pipeline by explicitly recording each transformation applied to the spectrum along with its parameters and execution history. This ensures that every processed spectrum can be traced back to its originating operations, supporting auditability and scientific transparency.
- **Open-source graphical user interface for standardized spectral processing.** We develop an open-source PyQt-based graphical application that exposes the full preprocessing pipeline through an interactive, step-by-step workflow. The GUI integrates calibration, correction, and spectral processing into a unified interface, lowering the barrier to adoption and reducing reliance on custom code.
- **Configuration-driven support for heterogeneous Raman instrumentation** TRaP introduces a configuration-driven mechanism that decouples preprocessing execution from instrument-specific acquisition settings. The system-level configuration determines which preprocessing stages (e.g., calibration and spectral response correction) are required for a given instrument, allowing a shared preprocessing pipeline to be applied across heterogeneous Raman systems (e.g., Clinical Fiber-optic Raman System, Commercial Raman Microscope, Commercial Portable System) while preserving system-dependent processing requirements.

## 2 System Architecture and Standardized Processing Workflow

### 2.1 Overall Software Design and Module Organization

TRaP is implemented as a desktop application that integrates a graphical user interface with a modular computational backend. The system architecture separates user interaction from numerical processing routines. The graphical interface manages user input, workflow navigation, and visualization, while the backend implements the spectral preprocessing algorithms, including calibration, spectral response correction, and preprocessing operations. This separation ensures that the computational routines remain reusable and can be executed consistently in both interactive and batch processing modes.

The application organizes Raman spectral preprocessing into a guided workflow consisting of five sequential stages: (0) configuration management, (1) X-axis calibration, (2) spectral response correction, (3) single-spectrum processing, and (4) batch processing. This ordering reflects the dependency structure of the underlying computations and mirrors the physical measurement chain of a dispersive Raman spectrometer. By enforcing this execution structure, TRaP reduces variability in preprocessing order across users and experimental sessions. The Fig. 2 shows the detailed pipeline for TRaP workflow.

**Fig 2:**
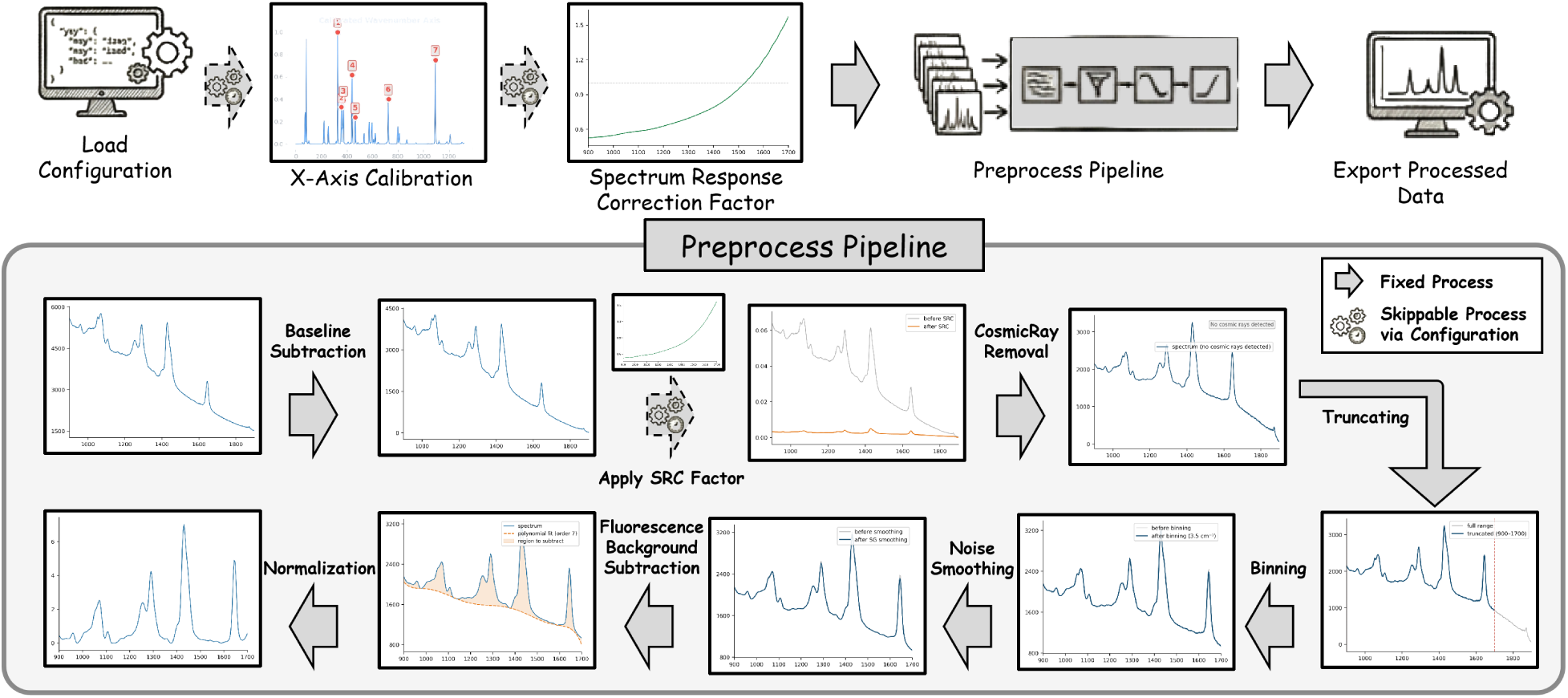
The TRaP spectral preprocessing pipeline and its role within the Raman analysis workflow. **Top:** TRaP is positioned within the end-to-end Raman workflow, including data acquisition, wavenumber calibration, spectral response correction, and downstream spectral processing. **Bottom:** The preprocessing pipeline implemented in TRaP consists of a sequence of configurable operations applied to input spectra. Starting from raw spectra, the pipeline performs baseline subtraction, optional spectral response correction (SRC), cosmic ray removal, truncation, binning, noise smoothing, fluorescence background subtraction, and normalization. Processing steps are categorized as either fixed or configurable, with selected stages (e.g., SRC) optionally skipped depending on instrument characteristics and configuration settings. This design enables consistent yet flexible preprocessing across datasets and acquisition systems.

The system maintains a structured data flow between processing stages. Configuration parameters defined at the beginning of the workflow determine the behavior of downstream steps. The calibration stage generates a wavelength mapping used by both correction and preprocessing modules, while the spectral response correction stage produces an instrument-specific correction factor that is subsequently applied during spectral preprocessing. This explicit parameter propagation ensures that all spectra processed within a session share a consistent set of calibration and correction parameters.

### 2.2 X-Axis Calibration (M1)

The X-axis calibration module as **M1** in Fig. 2 establishes a mapping from detector pixel positions to Raman shift values (cm^−1^). The procedure is implemented as a guided interactive workflow in which the user first selects reference peaks, then estimates the pixel-to-wavenumber mapping, and finally exports a reusable calibration record.

A subset of spectral lines is first selected from a built-in neon–argon emission reference library containing 52 emission lines spanning 9,677.994–12,947.060 cm^−1^ in absolute wavenumber. The user then uploads a measured neon–argon spectrum and marks the corresponding peaks on an interactive canvas. Peak positions are refined by applying a Savitzky–Golay first-derivative filter to identify peak spans and compute intensity centroids. A third-order polynomial is subsequently fitted to the paired pixel-position and absolute-wavenumber data by least-squares regression, yielding the mapping

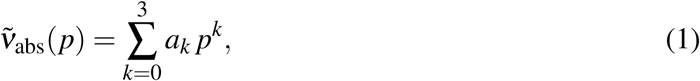

where *p* denotes the refined peak position and the coefficients {*a_k_*} are determined by the fit. The excitation laser wavelength required to convert absolute wavenumber to Raman shift may either be entered directly by the user or estimated from a measured acetaminophen reference spectrum. In the estimation pathway, Eq. (1) is evaluated at the pixel positions of selected acetaminophen peaks, and the laser wavelength is inferred as

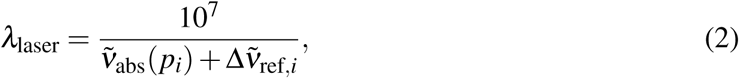

averaged over all selected peaks *i*, where Δ*ṽ*_ref_*_,i_* denotes the certified Raman shift of the *i*th acetaminophen reference peak drawn from a library of 22 certified peaks spanning 213.3– 3326.6 cm^−1^. The Raman shift axis is then computed as

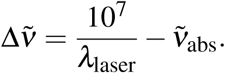

The calibration result is exported as a `.mat` file containing the wavenumber axis, laser wavelength, polynomial coefficients, calibration date and time, and the indices of the selected reference peaks. This output provides a documented and reloadable calibration record that can be reused in subsequent preprocessing sessions.

### 2.3 Spectral Response Correction (M2)

The spectral response correction module as **M2** in Fig. 2 compensates for wavelength-dependent sensitivity in the detector and optical path by computing a multiplicative correction factor *C*(*ṽ*) applied to each raw spectrum. The purpose of this module is not to introduce a new correction model, but to standardize several commonly used correction pathways within a single controlled interface. Three correction modes are currently supported.

#### White-light correction

A measured spectrum of a calibrated broadband light source is smoothed and converted to scattered wavelength in nanometres via *λ* = 10^7^*/*(10^7^*/λ*_laser_ −Δ*ṽ*). A polynomial is fitted to the manufacturer-provided reference radiance table and evaluated at the calibrated wavelength axis to obtain the true spectral profile. Both profiles are normalized at a common centre wavelength, and the correction factor is computed as their ratio.

#### NIST standard reference material correction

A measured spectrum of a NIST-certified fluorescent glass standard is used as input. The true spectral profile is evaluated using the certified fifth-order polynomial coefficients specified in the NIST standard reference material documentation. The correction factor is derived as the ratio of the normalized true and measured SRM profiles, followed by smoothing.

#### Pre-computed correction factor

A previously computed correction factor may be loaded directly from a text or spreadsheet file, bypassing the computation stage. This option is intended for instrument configurations for which a valid correction factor has already been established in a prior session.

### 2.4 Spectrum Preprocessing Pipeline (M3)

The core preprocessing pipeline shown is invoked identically from both the interactive single-spectrum interface and the batch processing module. As shown in lower panel of Fig. 2, the preprocessing workflow consists of eight computational operations (**P1–P8**)applied in a fixed order within the user interface. This ordering is enforced to ensure deterministic and reproducible preprocessing across exploratory and production use.

#### P1. Baseline subtraction

The minimum intensity value is subtracted from the raw spectrum, shifting the electronic baseline to zero:

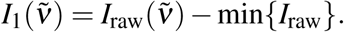

#### P2. Apply spectral response correction factor

The correction factor *C*(*ṽ*) is applied multiplicatively. For data formats in which the calibrated wavenumber axis is already supplied and the TRaP workflow does not require an externally provided correction file, this step is bypassed.

#### P3. Cosmic ray handling

A reserved processing stage is included to maintain a consistent pipeline structure and allow future integration of cosmic-ray removal algorithms. In the current implementation, this step does not modify the spectrum.

#### P4. Spectral truncating

The spectrum and wavenumber axis are cropped to a user-specified range [*ṽ*_start_, *ṽ*_stop_], with a default of 900–1700 cm^−1^ corresponding to the Raman fingerprint region.

#### P5. Spectral binning

The truncated spectrum is resampled onto a uniform wavenumber grid with spacing Δ*b* (default 3.5 cm^−1^) by averaging all data points within each bin interval:

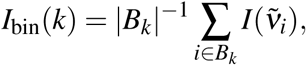

where *B_k_* denotes the set of indices within the *k*-th bin.

#### P6. Noise smoothing

One of three smoothing filters—Savitzky–Golay, moving average, or median filtering—may be applied. This step can be skipped if smoothing is not required.

#### P7. Fluorescence background subtraction

A broad fluorescence baseline is estimated using an iterative asymmetric polynomial fitting procedure. At each iteration, a polynomial of user-specified order is fitted and the spectrum is updated by taking the pointwise minimum of the fit and current values. The final baseline is then subtracted from the spectrum.

#### P8. Normalization

The processed spectrum is divided by its mean intensity,

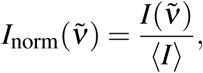

yielding a standardized spectral profile. The processed spectrum is exported as a two-column file containing the wavenumber axis and normalized intensity. Alternative normalization strategies, including area-based normalization and maximum-intensity normalization, are also supported within the same framework. The processed spectrum is exported as a two-column file containing the wavenumber axis and normalized intensity.

## 3 Configuration-Driven Processing for Heterogeneous Systems and Reproducible Workflows

TRaP addresses two central challenges in Raman spectral analysis: the heterogeneity of acquisition systems and the reproducibility of preprocessing workflows. This section describes the configuration-driven system architecture that enables spectral processing across heterogeneous instruments while ensuring reproducible workflow execution.

### 3.1 System-Level Configuration for Heterogeneous Instruments

Raman spectra acquired from different instruments exhibit substantial heterogeneity in acquisition configuration, calibration requirements, and data representation. Variations in excitation wavelength, detector characteristics, optical setup, and vendor-specific export formats result in differences in whether calibration or spectral response correction is required prior to preprocessing.

As illustrated in Fig. 3, TRaP handles this heterogeneity through a system-level configuration file (`config.json`). At the beginning of the workflow, the user specifies the instrument type and associated acquisition settings. This configuration encodes system-specific requirements and determines which preprocessing stages must be executed.

**Fig 3:**
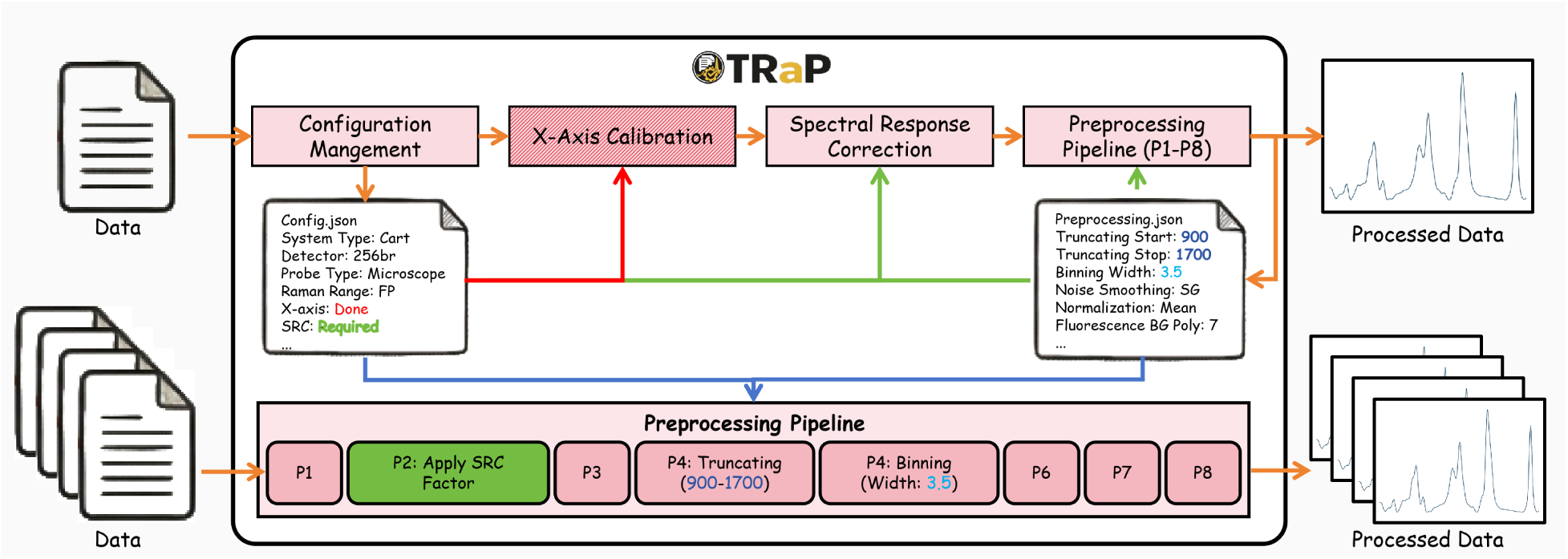
Overview of the TRaP framework for configuration-driven Raman spectral preprocessing. The upper panel illustrates the end-to-end workflow. Instrument- and experiment-specific parameters are encoded in configuration files (e.g., `config.json` and `preprocessing.json`), enabling explicit control over processing behavior. The lower panel details the deterministic preprocessing sequence, in which each operation is executed in a fixed order with predefined parameters (e.g., truncating over 900–1700 cm^−1^, binning with a width of 3.5, and application of the SRC factor due to Clinical Fiber-optic Raman system configuration). All processing steps and parameters are explicitly recorded, ensuring full provenance tracking and transparency.

In particular, the configuration controls the activation of X-axis calibration and spectral response correction (SRC). For instruments that provide calibrated spectra (e.g., Commercial Raman Microscope systems), these steps are automatically bypassed. For instruments that provide intensity-only measurements, external calibration and correction inputs are required and incorporated into the workflow.

Importantly, this configuration does not alter the preprocessing pipeline itself. Instead, it defines the execution path through the workflow by enabling or disabling specific stages. This design allows heterogeneous Raman systems to be processed within a unified framework while preserving instrument-dependent processing requirements.

### 3.2 Pipeline-Level Parameterization for Reproducible Processing

Given the execution path defined by the system-level configuration, TRaP ensures reproducible preprocessing through a parameter-driven pipeline specification. As shown in Fig. 3, all preprocessing operations (P1–P8) are governed by a structured parameter file (`preprocessing.json`), which defines both the sequence of operations and their associated parameters.

Each preprocessing step—including truncation range, binning width, smoothing method, and background subtraction—is explicitly specified within this configuration. Unlike conventional preprocessing workflows implemented as ad hoc scripts, TRaP encodes the entire pipeline in a declarative and portable format, eliminating ambiguity in algorithm selection, parameter values, and execution order.

The preprocessing pipeline is executed deterministically: all operations are applied in a fixed order with no hidden defaults, and the same implementation is shared across both interactive and batch processing modes. As a result, identical input data processed under the same configuration produce identical outputs.

Furthermore, all parameters and processing steps are recorded, enabling full provenance tracking and transparent inspection of the preprocessing workflow. Because the configuration is portable, preprocessing procedures can be shared and re-executed across users, datasets, and computational environments without modification.

## 4 Experiments

### 4.1 Reproduction Experiments

To demonstrate how TRaP supports reproducible workflow sharing, we compared spectra generated by original instrument-specific preprocessing pipelines with spectra produced by TRaP configured to replicate the same procedures. For each Raman platform, spectra were processed using two strategies: (1) the original instrument-specific pipeline used in prior practice, and (2) a TRaP workflow configured with the same sequence of processing operations and parameter settings. In TRaP, these workflow definitions are stored as reusable configuration files rather than embedded in scripts or reconstructed manually by users. As shown in Fig. 4, spectra processed by TRaP closely match those produced by the native pipelines in peak position, peak shape, and relative intensity structure. This agreement illustrates that existing processing procedures can be reproduced while being converted into explicit and shareable configuration records. By representing processing protocols as configuration files rather than implicit software settings or custom scripts, TRaP enables Raman spectral preprocessing workflows to be distributed, reused, and reproduced across users and computational environments. Other researchers can reproduce the same processing procedure by loading the shared configuration together with the corresponding spectral data.

**Fig 4:**
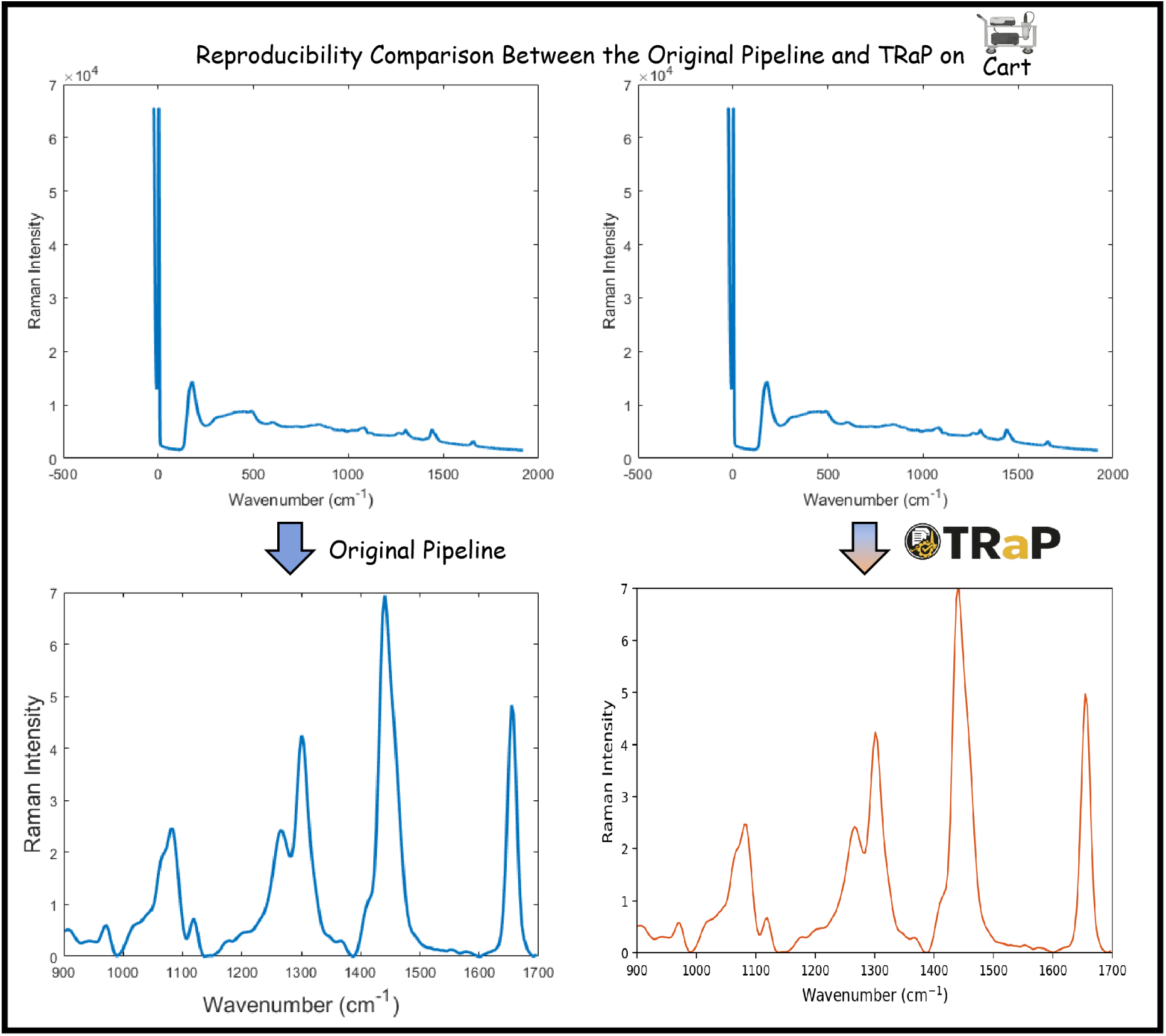
Reproduction of instrument-specific Raman preprocessing workflows using TRaP. For each Raman platform, spectra processed using the original instrument-specific pipeline are compared with spectra processed using TRaP configured to replicate the same procedure. The close agreement in peak positions, peak shapes, and relative intensity patterns demonstrates that legacy preprocessing workflows can be reproduced and executed within a unified configuration-driven framework.

### 4.2 Detailed User Case

Figure 5 illustrates the graphical workflow implemented in TRaP. The system organizes Raman spectral preprocessing into a configuration-driven workflow within a unified interface, where instrument metadata (e.g., system type, excitation wavelength, detector, and probe configuration) define subsequent calibration and processing behavior. Calibration steps—including X-axis calibration and spectral response correction—are applied conditionally based on the selected configuration, while systems with pre-calibrated spectral axes can bypass these steps automatically.

**Fig 5:**
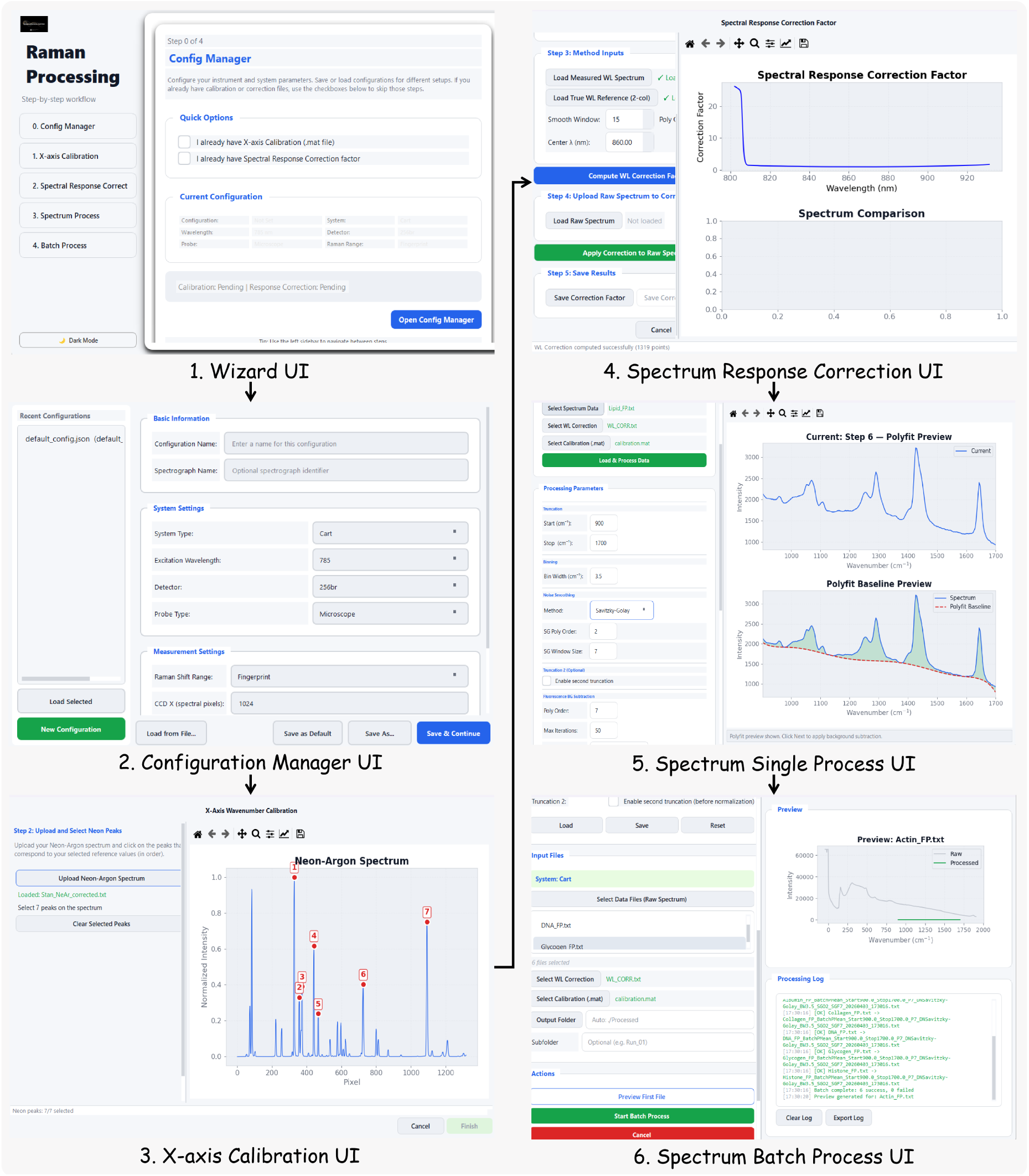
Overview of the TRaP application workflow. The graphical interface guides users through the complete Raman processing pipeline, including (1) workflow wizard entry, (2) instrument configuration management, (3) X-axis calibration using reference spectra, (4) spectral response correction, (5) single-spectrum processing, and (6) batch processing of multiple spectra.

Cross-system preprocessing is enabled through reusable configuration files that encapsulate instrument-specific parameters, allowing spectra from heterogeneous Raman systems to be processed under a shared workflow. Processing parameters are consistently applied across datasets, ensuring alignment to a common wavenumber grid and output format. Both interactive and batch execution modes invoke the same computational pipeline, guaranteeing deterministic and reproducible preprocessing under identical configurations. Detailed step-by-step usage instructions are provided in Appendix 6.

## 5 Discussion

### 5.1 Summary

This paper presents TRaP, an open-source framework for configuration-driven and reproducible Raman spectral preprocessing across heterogeneous acquisition systems. TRaP separates system-level execution control from pipeline-level parameter specification through structured configuration files. Instrument-specific differences are encoded in system configurations, which determine the activation of preprocessing stages such as calibration and spectral response correction, while a shared preprocessing pipeline is applied across all supported instruments. Preprocessing parameters are explicitly defined and consistently applied, and the same pipeline implementation is used for both interactive and batch execution modes. This design ensures deterministic behavior, such that identical input data processed under the same configuration yield identical outputs. All processing steps and parameters are recorded, enabling full provenance tracking and transparent inspection of the preprocessing workflow. Together, these design choices establish a reproducible and transparent preprocessing framework for Raman spectroscopy.

### 5.2 Strengths and Limitations

TRaP provides a configuration-driven framework that separates system-dependent execution control from pipeline-level parameter specification, enabling consistent preprocessing across heterogeneous Raman acquisition systems. By encoding instrument-specific requirements in system configurations, the framework determines the activation of preprocessing stages (e.g., calibration and spectral response correction) without modifying the underlying pipeline. This design allows a shared preprocessing pipeline to be applied across different instruments while preserving system-dependent processing requirements.

Reproducibility is further supported through explicit parameterization of the preprocessing workflow. All operations and associated parameters are defined in configuration files, eliminating ambiguity in algorithm selection, parameter values, and execution order. The use of a single pipeline implementation across both interactive and batch modes ensures deterministic behavior, such that identical input data processed under the same configuration produce identical outputs. In addition, all processing steps and parameters are recorded, enabling full provenance tracking and transparent inspection of the preprocessing workflow.

Despite these strengths, TRaP has several limitations. The framework does not address variability introduced during data acquisition, such as differences in sample preparation, laser power, or environmental conditions, which may affect spectral quality independently of preprocessing. The configuration system relies on predefined instrument categories and constraints, which may limit flexibility when adapting to novel or highly customized acquisition setups. Furthermore, TRaP focuses exclusively on preprocessing and does not extend to downstream analytical tasks such as peak interpretation, classification, or quantitative modeling; therefore, reproducibility is ensured at the preprocessing stage but not necessarily across the full analytical pipeline.

### 5.3 Future Work

Future work will extend TRaP in several directions. First, support for additional Raman systems and more flexible configuration schemas will be developed to accommodate a broader range of experimental setups. Second, integration with downstream analysis modules, such as peak detection, spectral unmixing, and machine learning-based classification, will enable end-to-end reproducible Raman workflows. In addition, systematic evaluation of preprocessing sensitivity to parameter variation and upstream acquisition variability will be conducted to better characterize the robustness of the pipeline. Finally, the incorporation of standardized benchmark datasets and shared configuration repositories may further promote reproducibility and facilitate community-driven development.

## 6 Conclusion

This paper presents TRaP, an open-source framework for reproducible Raman spectral preprocessing across heterogeneous acquisition systems. TRaP integrates instrument configuration management, X-axis calibration, spectral response correction, and a standardized preprocessing pipeline into a unified, configuration-driven workflow. Rather than treating preprocessing steps as independent utilities, the framework formalizes spectral preprocessing as a deterministic execution pipeline whose behavior is fully specified through shareable configuration files.

TRaP formalizes preprocessing as a configuration-driven and deterministic execution process. All processing steps, parameter values, and calibration outputs are explicitly encoded in reusable configuration files, eliminating ambiguity in workflow definition. A single pipeline implementation is used across both interactive and batch execution modes, ensuring that identical inputs and configurations produce identical outputs. This design establishes a transparent, reproducible, and transferable preprocessing paradigm for Raman spectroscopy. The framework further supports cross-system comparability by decoupling instrument-specific variability from preprocessing logic, allowing consistent analysis across heterogeneous datasets. At the same time, TRaP focuses specifically on preprocessing and does not address variability introduced during data acquisition or downstream analytical tasks, which remain important factors in end-to-end reproducibility.

TRaP is not intended to replace specialized Raman analysis platforms, but to provide a structured and instrument-agnostic preprocessing framework for Raman spectra. Future work will extend TRaP toward broader system support, more flexible configuration schemas, and integration with downstream analytical modules such as peak detection and machine learning-based modeling. In addition, systematic evaluation of preprocessing robustness and the development of shared benchmark datasets and configuration repositories will further strengthen reproducibility and facilitate community adoption.

## Acknowledgments

This research was supported by NIH R01DK135597 (Huo), DoD HT9425-23-1-0003 (HCY), and KPMP Glue Grant. This work was also supported by Vanderbilt Seed Success Grant, Vanderbilt Discovery Grant, and VISE Seed Grant. This project was supported by The Leona M. and Harry B. Helmsley Charitable Trust grant G-1903-03793 and G-2103-05128. This research was also supported by NIH grants R01EB033385, R01DK132338, REB017230, R01MH125931, and NSF 2040462. We extend gratitude to NVIDIA for their support by means of the NVIDIA hardware grant. This work was also supported by NSF NAIRR Pilot Award NAIRR240055.

During the preparation of this work the author used ChatGPT in order to improve the grammar and clarity of the writing. In addition, Claude Code was used to assist with repository management, debugging, and code refinement. All functions and features were subsequently reviewed, revised, and validated through human testing. After using this tool/service, the author reviewed and edited the content as needed and takes full responsibility for the content of the manuscript.

## Appendix A: Step-by-Step Workflow for TRaP

This appendix provides a visual walkthrough of the complete TRaP workflow, from initial configuration through batch export using representative Raman spectroscopy data consistent with the protocol described by Haugen et al.^19^ The workflow consists of four main stages: system configuration, X-axis calibration, spectral response correction, and spectral processing (Spectrum Processing or Spectrum Batch Processing).

### A.1 Launching TRaP and System Configuration

Upon launching TRaP, the main wizard window is presented (Figure 6). The wizard guides the user through Steps 0–4. Before proceeding, the system configuration must be completed by clicking

#### Open Config Manager

The Config Manager dialog (Figure 7) organises settings into three groups:

**Fig 6:**
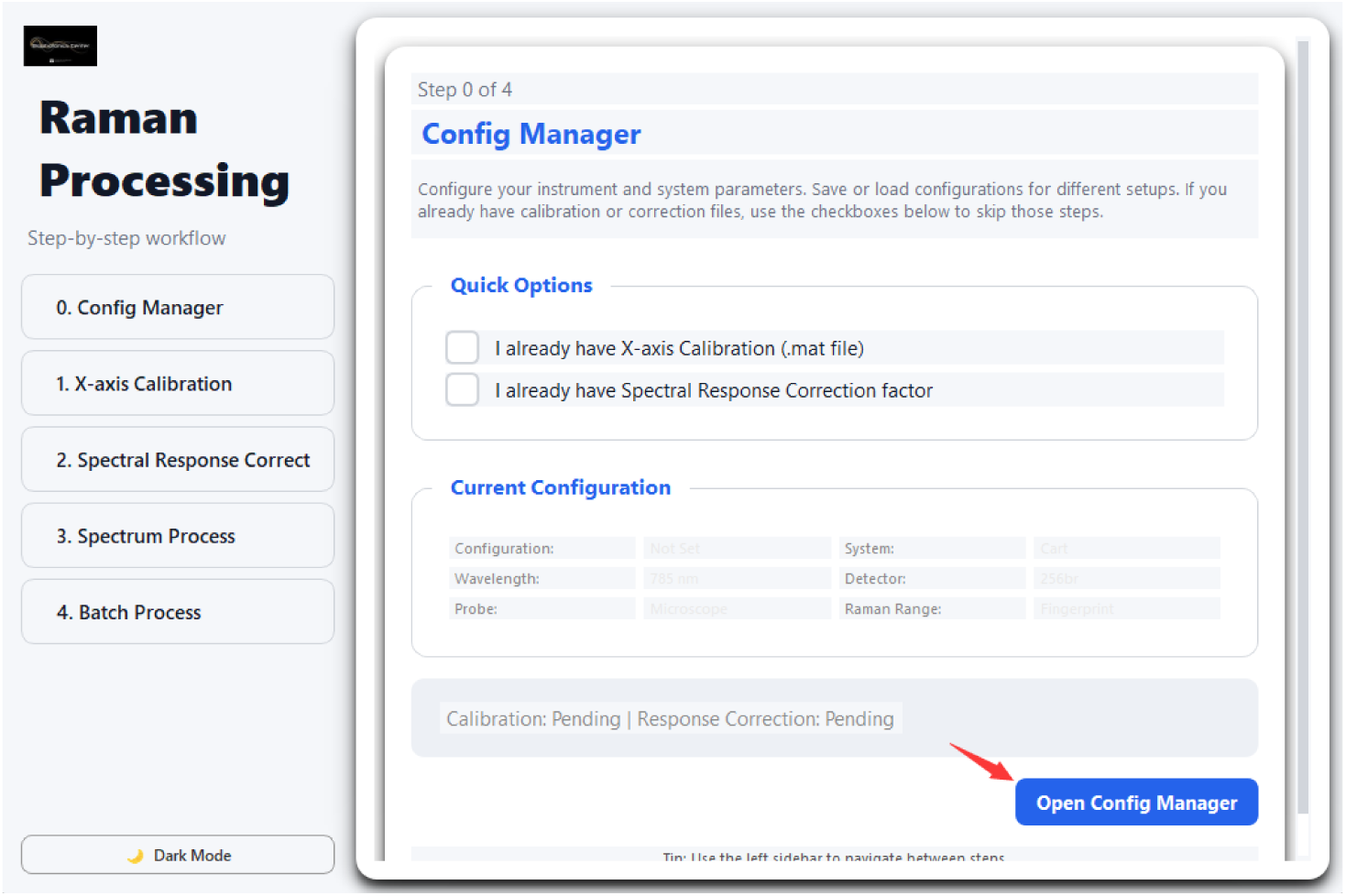
TRaP main wizard screen (Step 0 of 4). The *Config Manager* button opens the configuration dialog before any processing begins.

- **Basic Info** — operator name and experiment label.
- **System Settings** — instrument system (e.g. Clinical Fiber-optic Raman System, Commercial Raman Microscope), excitation wavelength, detector model, and probe type. Selecting a detector automatically pre-fills the CCD pixel dimensions.
- **Measurement Settings** — spectral range (Start/Stop in cm^−1^) and CCD pixel counts (X: spectral, Y: spatial), which may be edited manually after auto-fill.

Click **Save & Close** to persist the configuration to `defaultconfig.json`.

**Fig 7:**
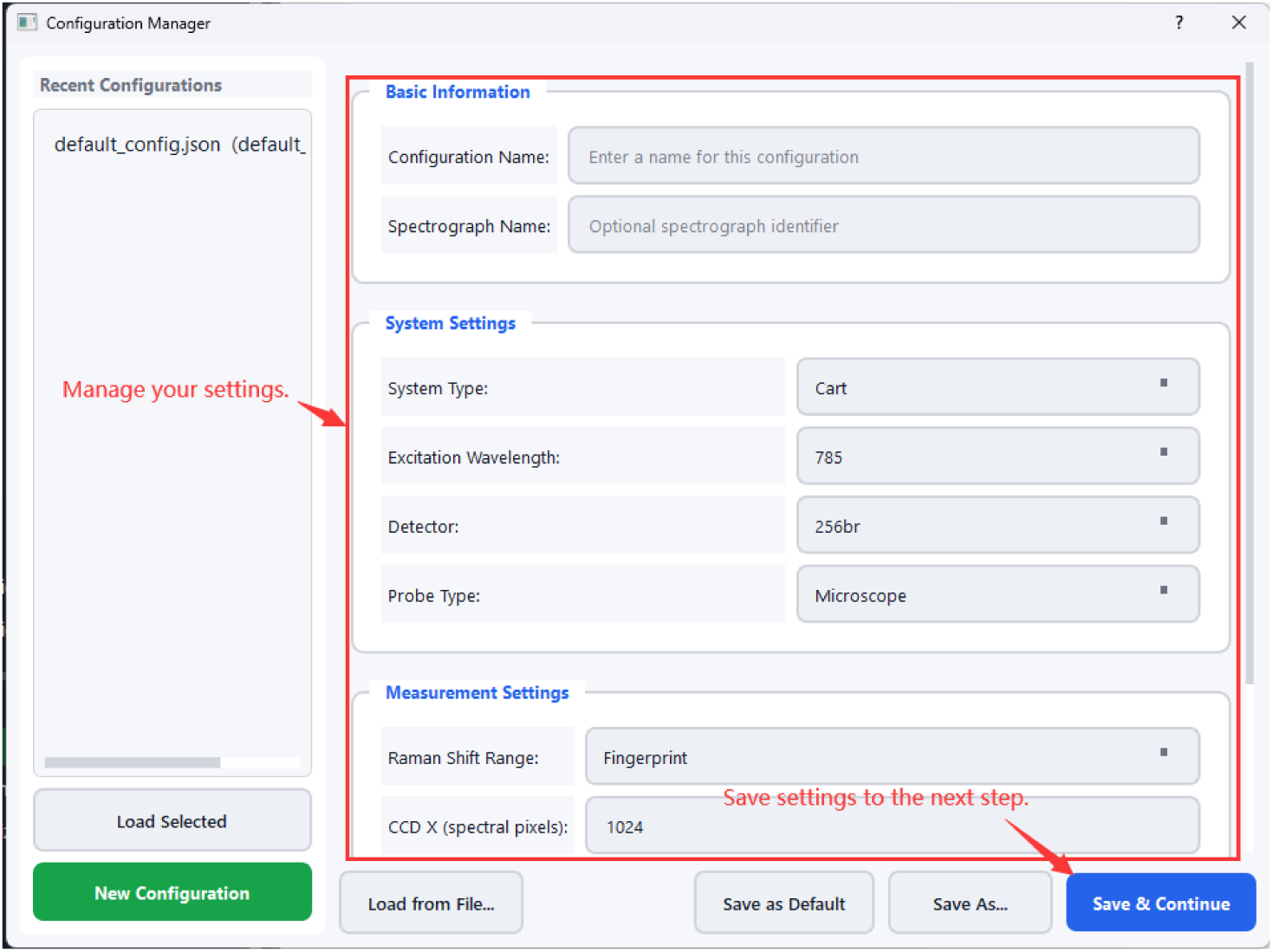
Config Manager dialog. System Settings (System, Wavelength, Detector, Probe) and Measurement Settings (spectral range, CCD dimensions) are configured here. Selecting a detector pre-fills CCD X and Y values.

### A.2 X-axis Calibration

The X-axis calibration step corresponds to the wavelength calibration procedure described in standard Raman spectroscopy protocols, where known emission lines are used to establish a mapping between detector pixels and Raman shift. Step 1 of the wizard performs X-axis (wavenumber) calibration using a Neon–Argon lamp spectrum. Click Start X-axis Calibration (Figure 8) to open the calibration dialog.

**Fig 8:**
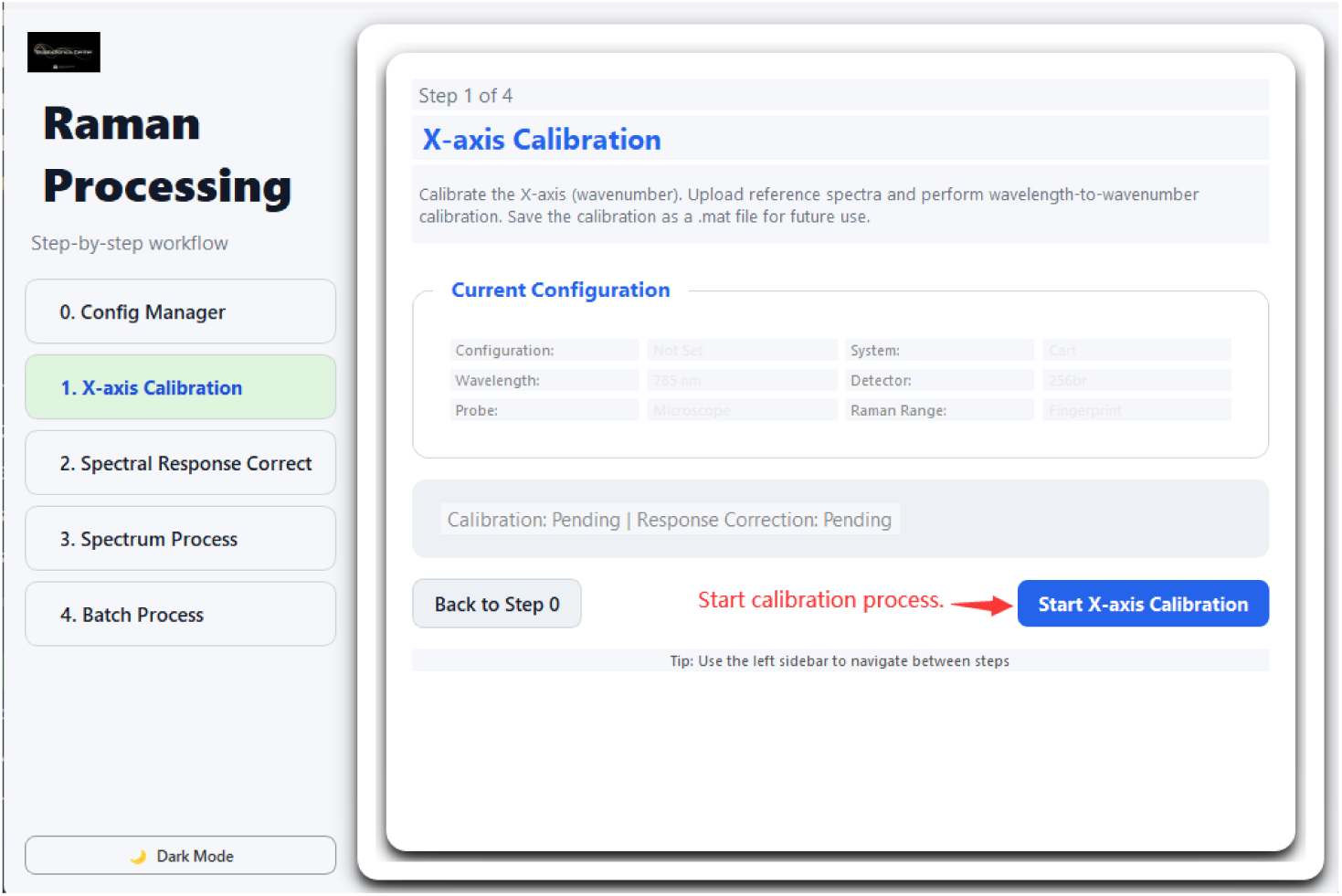
Wizard Step 1: X-axis Calibration entry point. Clicking *Start X-axis Calibration* opens the calibration dialog.

**Step 1 — Select reference peaks.** In the *Neon Reference Library* panel, select the Ne emission lines that will be used for fitting. It is recommended to select 6–8 well-isolated peaks spanning the full spectral range (Figure 9).

**Step 2 — Upload the Ne–Ar spectrum.** Click Upload Ne-Ar Spectrum to load the raw lamp spectrum acquired with the instrument (Figure 10).

**Step 3 — Mark peaks on the spectrum.** Once the spectrum is loaded, click on each peak position in the plot that corresponds to the selected reference lines. The annotated positions are listed in the right panel (Figure 11).

**Step 4 — Enter laser wavelength.** The laser wavelength is required to convert pixel positions to Raman shift (cm^−1^). Option A (recommended) is to enter the value directly (e.g. 785.142nm).

**Fig 9:**
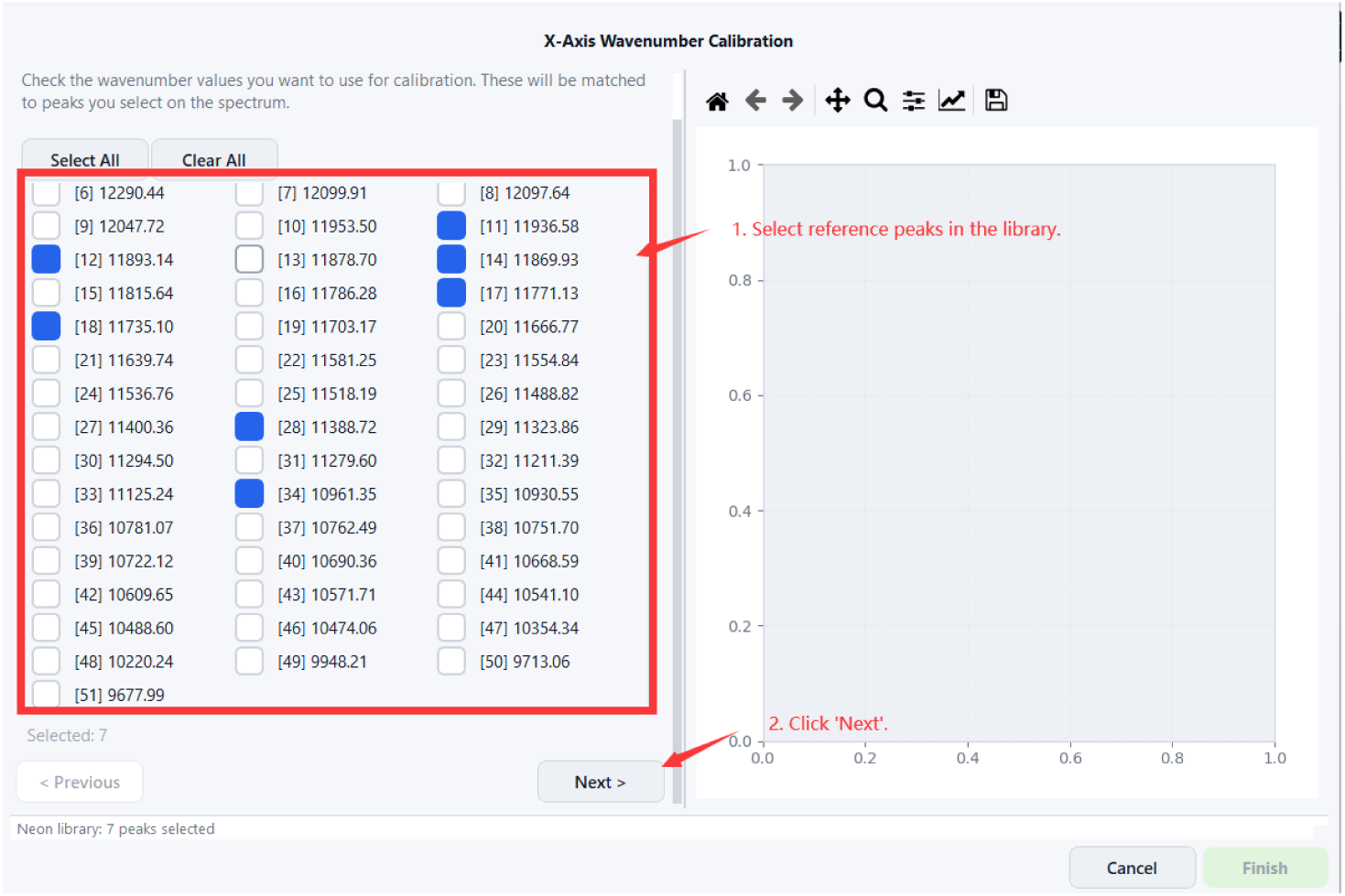
Calibration Step 1: selecting Neon reference peaks from the built-in library. Seven peaks are selected (shown in blue).

**Fig 10:**
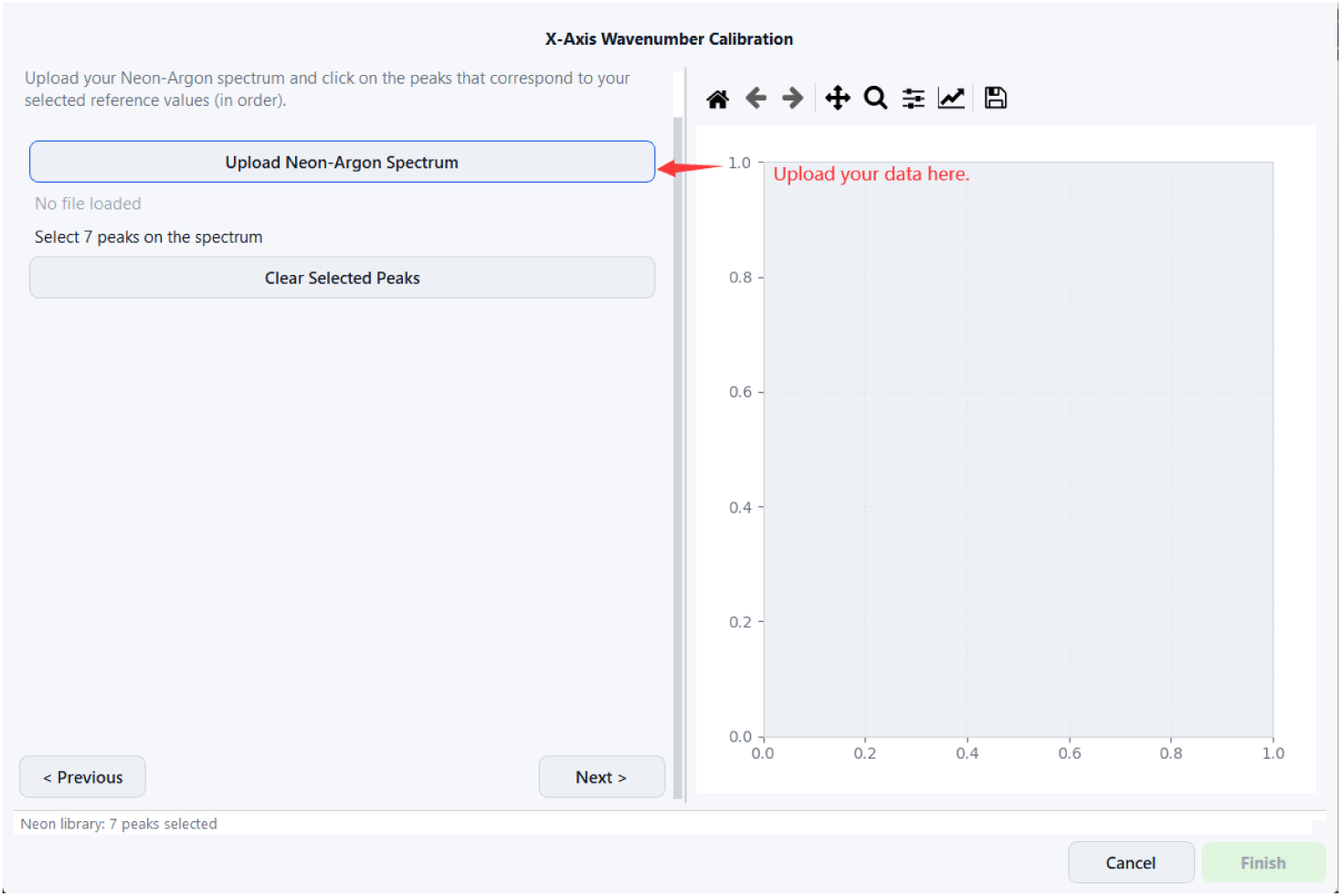
Calibration Step 2: uploading the measured Neon–Argon lamp spectrum file.

**Fig 11:**
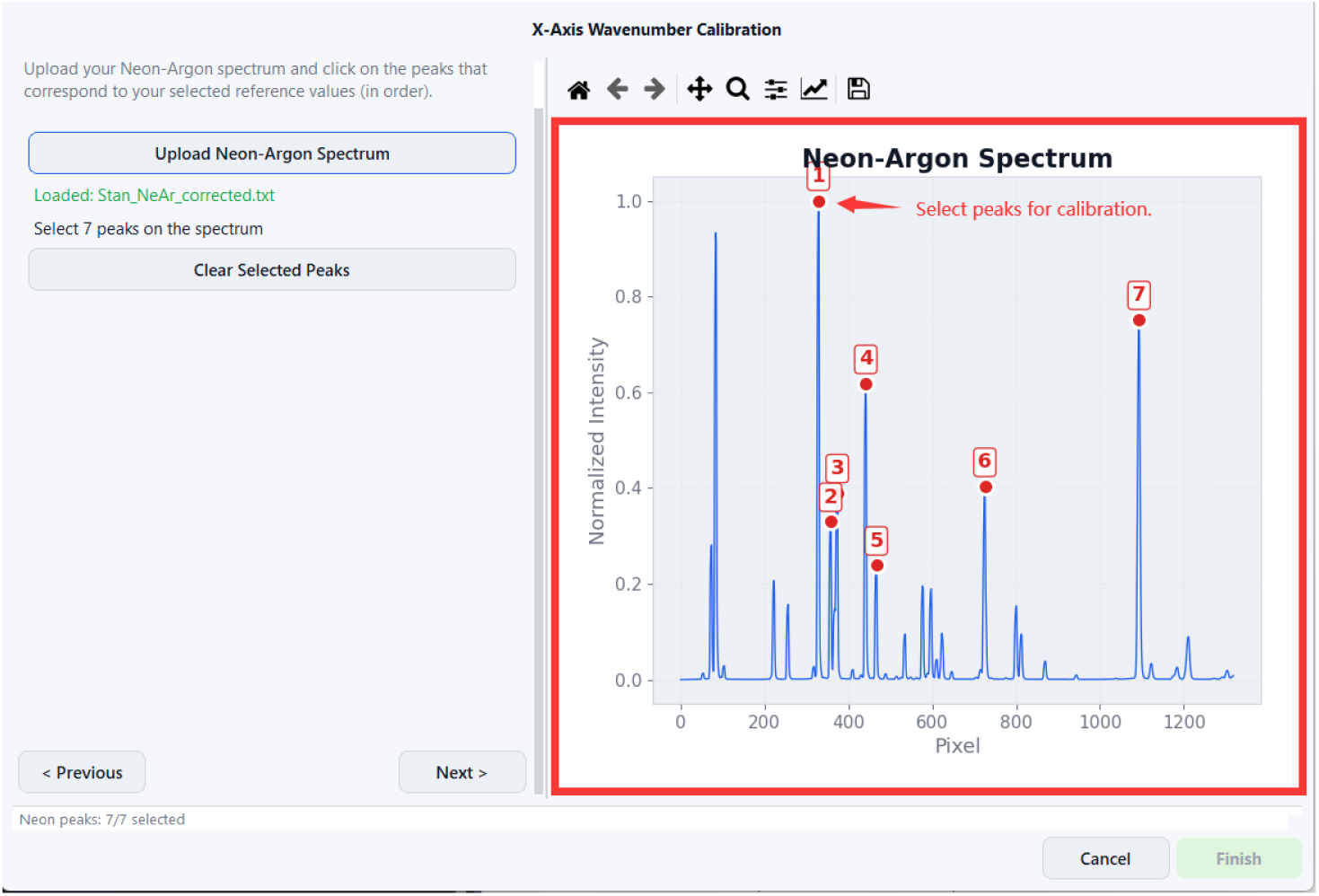
Calibration Step 3: interactively clicking peak positions on the loaded Ne–Ar spectrum. Marked positions appear in the right-hand table.

Option B uses Acetaminophen peaks as an internal standard (Figure 12).

**Fig 12:**
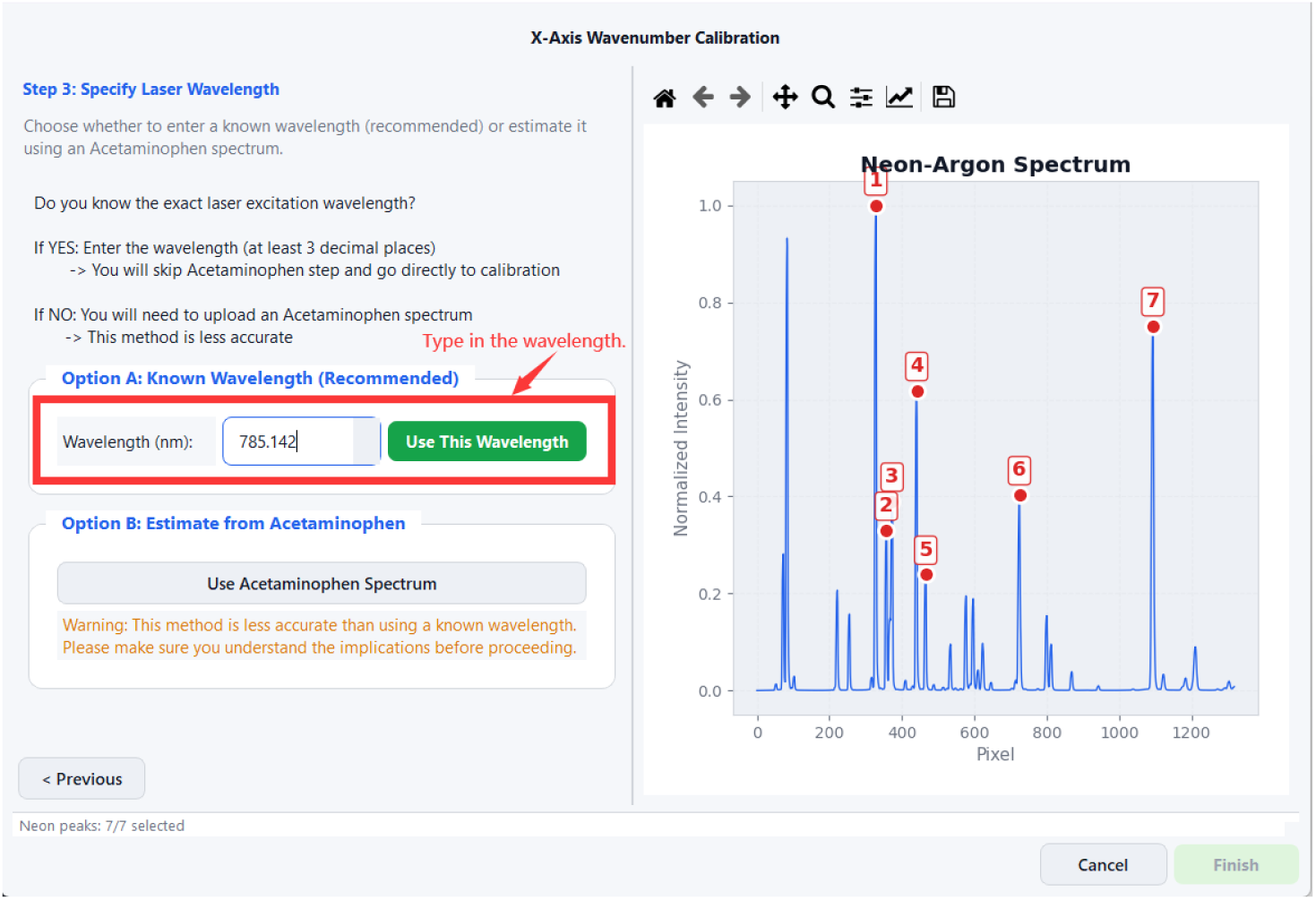
Calibration Step 4: entering the excitation laser wavelength. Direct entry (Option A, 785.142nm) is the recommended approach.

**Step 5 — Run calibration and save. Click Run Calibration.** TRaP fits a polynomial mapping from pixel index to wavenumber. Upon success, the mean absolute error (MAE) is reported (e.g. 0.0561cm^−1^) and the calibration is saved as a `.mat` file for use in subsequent steps (Figure 13).

**Fig 13:**
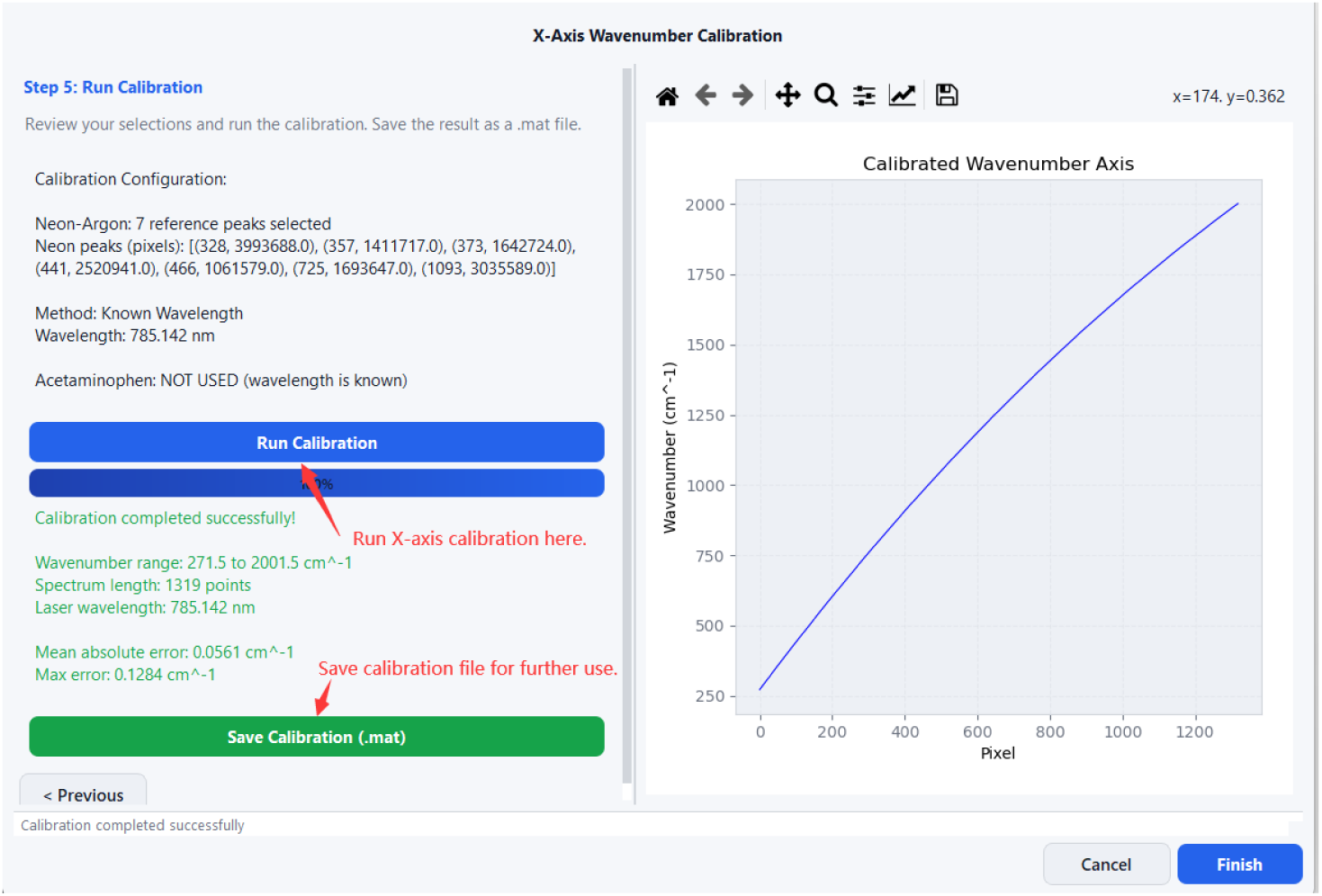
Calibration Step 5: successful calibration result. The calibration is saved as a `.mat` file.

### A.3 Spectral Response Correction

Step 2 of the wizard corrects for the wavelength-dependent response of the detector and optics. Spectral response correction follows the common practice of normalizing the system response using a broadband reference source, ensuring that measured spectra are comparable across systems. Click **Open Response Correction** (Figure 14) to open the Spectral Response Correction dialog. In the SRCF dialog (Figure 15), select the correction method (*White Light*), load the calibration `.mat` file from Step 1, then load both the measured white-light spectrum and the true (reference) white-light spectrum.

**Fig 14:**
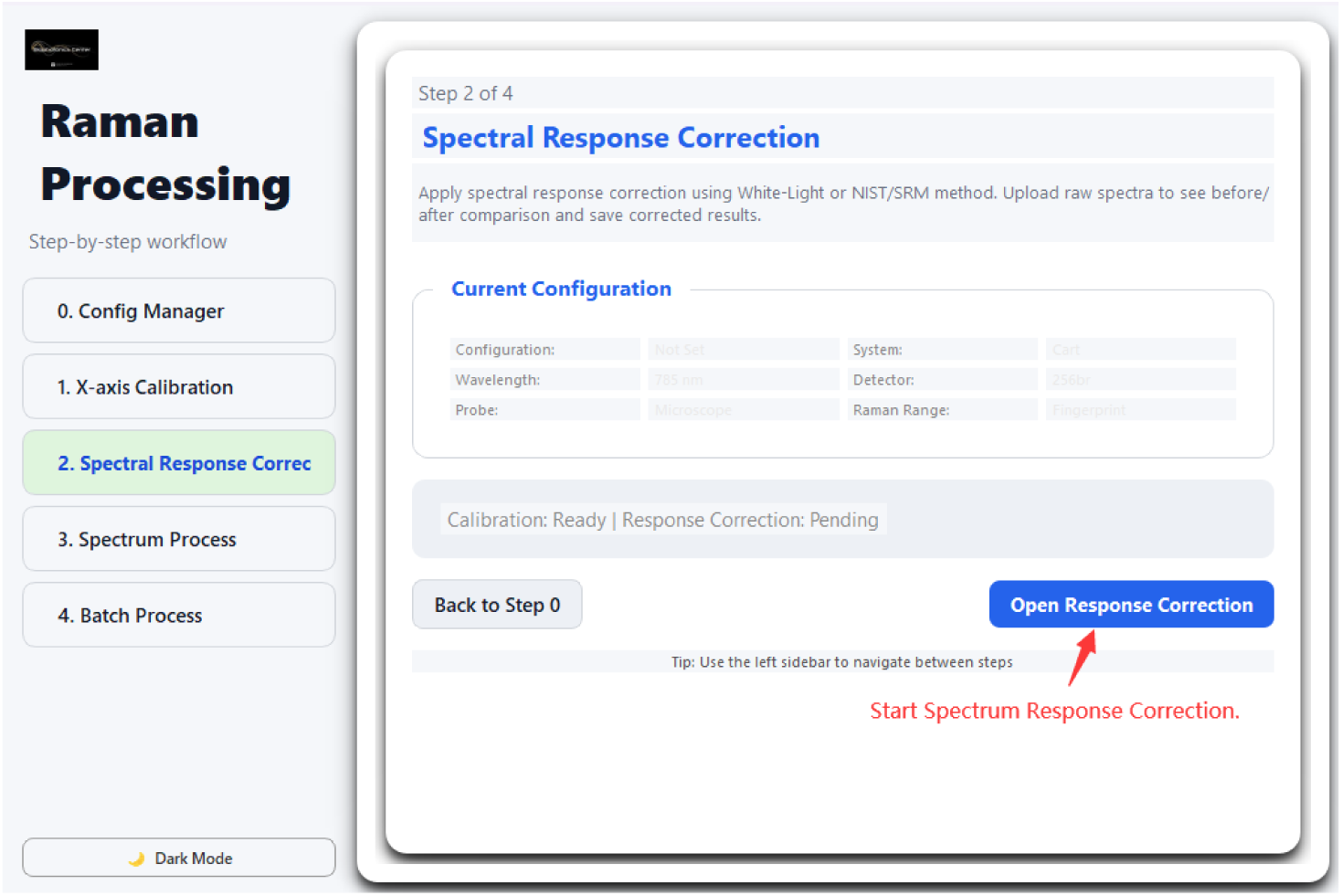
Wizard Step 2: Spectral Response Correction entry point.

**Fig 15:**
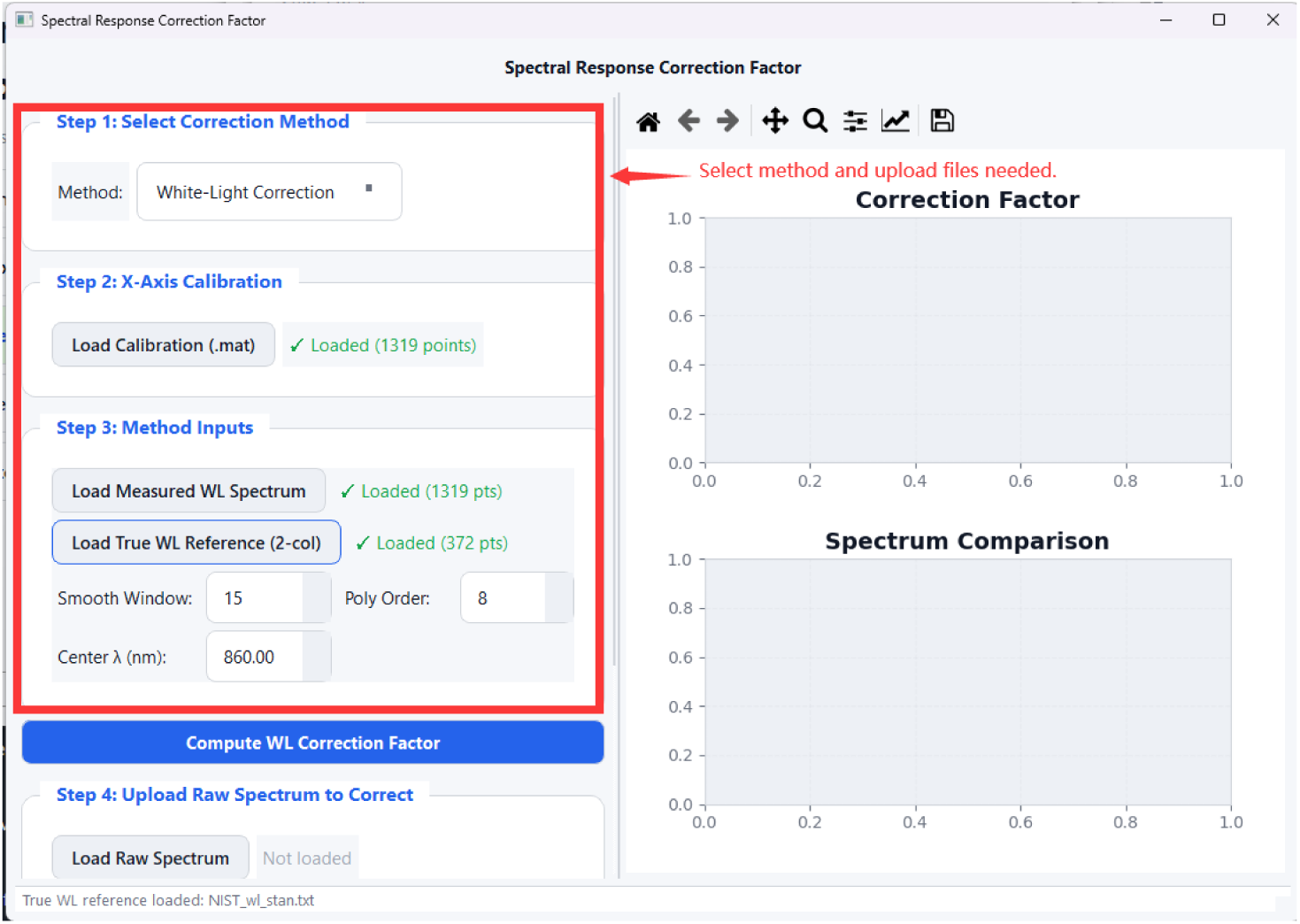
SRCF dialog: selecting the White Light method, loading the calibration file, and providing the measured and reference white-light spectra.

Click **Compute WL Correction Factor.** The computed correction curve is displayed in the plot.

Click **Save Correction Factor** to export the factor as `WLCORR.txt`, which will be used in all subsequent spectral processing (Figure 16).

**Fig 16:**
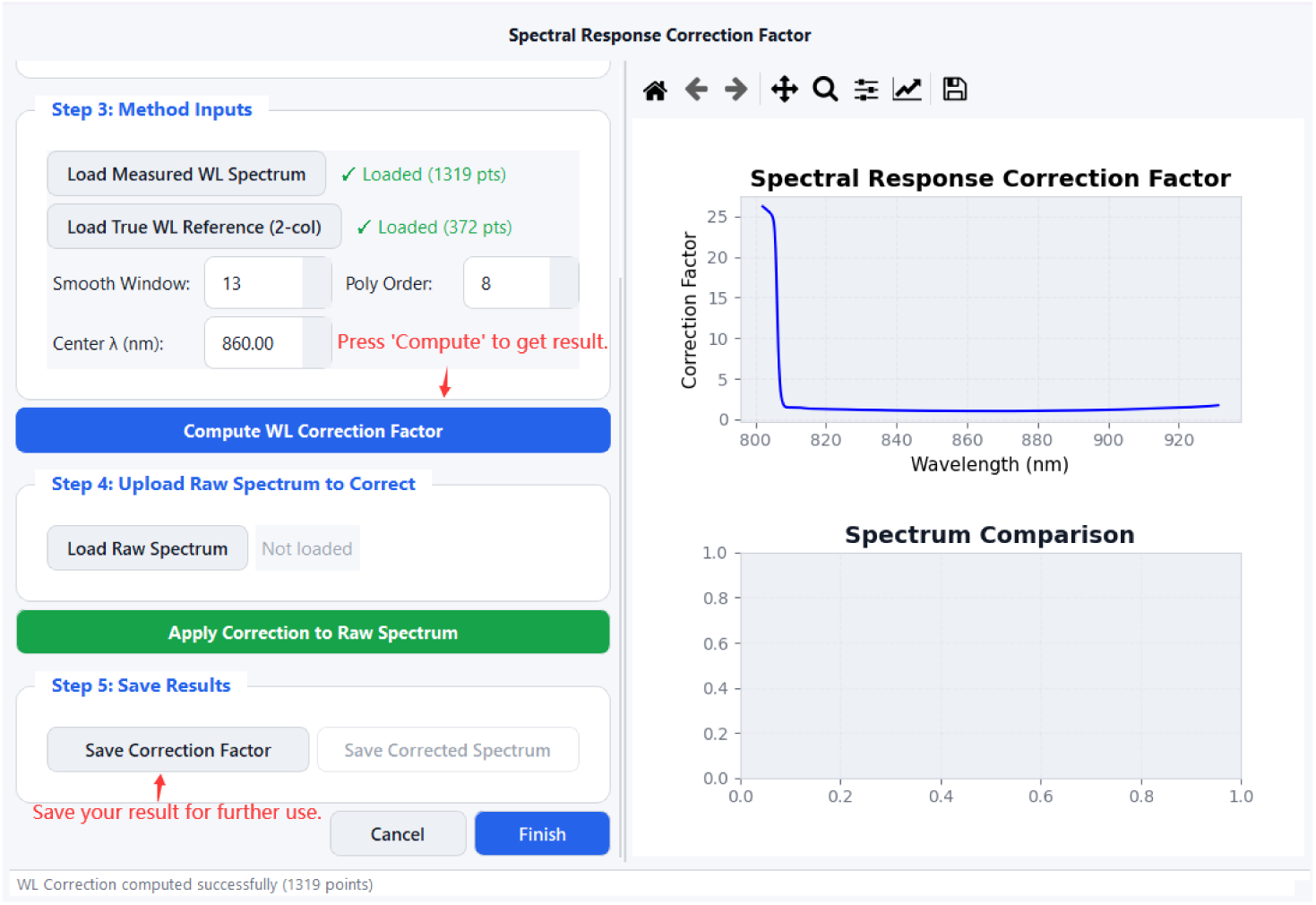
SRCF result: the computed wavelength correction factor curve is displayed and saved as WLCORR.txt.

### A.4 Spectrum Processing

Steps 3–4 of the wizard perform per-spectrum processing. The spectral processing stage in TRaP implements the core preprocessing steps widely adopted in Raman spectroscopy, including truncation, binning, fluorescence background subtraction, noise smoothing, and normalization. The left panel is divided into **Input Files** and **Processing Parameters** sections.

**Input Files** (Figure 17): select raw spectrum data files, the WL correction factor (`WLCORR.txt`) from Step 2, and the calibration file (`.mat`) from Step 1. Click **Load & Process Data** to begin processing.

**Processing Parameters** (Figure 18): the pipeline parameters are organised by step:

- **Truncation** — Start and Stop wavenumber (cm^−1^) to define the spectral window of interest.
- **Binning** — bin width in cm^−1^ (default 3.5).

**Fig 17:**
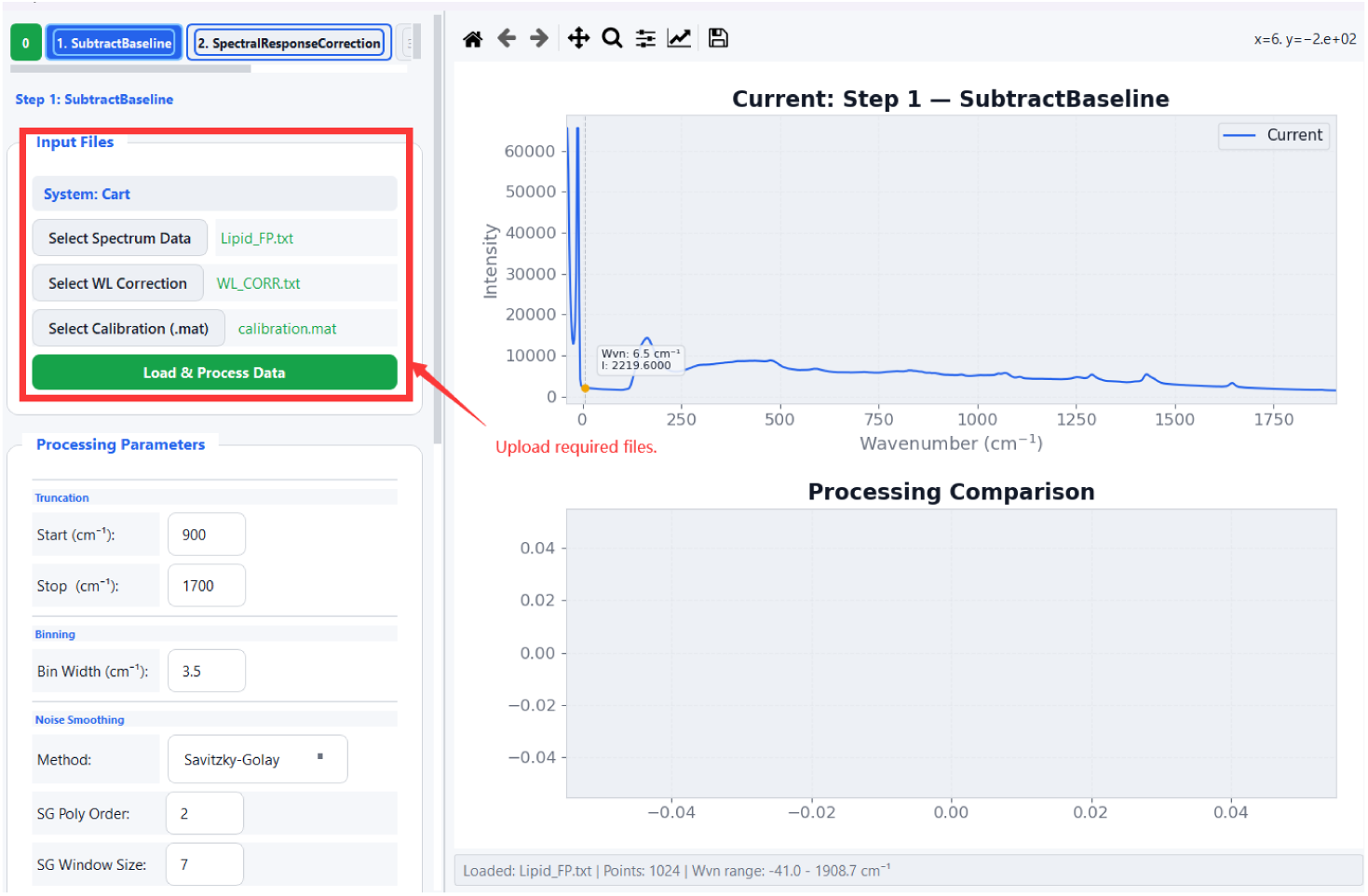
Spectrum Processing: Input Files panel showing data file selection, WL correction, and calibration inputs, with the *Load & Process Data* button.

- **Fluorescence Background Subtraction (FBS)** — polynomial order and maximum iterations for the iterative baseline fit.
- **Normalization** — method selection (Mean, Max, or Area).
- **Noise Smoothing** — method (Savitzky–Golay, Moving Average, or Median), with method-specific parameters (SG polynomial order and frame size, or window size) shown conditionally.

Configurations can be saved to or loaded from a JSON file using **Save/Load Config**.

After processing, the result is displayed in the dual-plot view (Figure 19). The upper plot shows the spectrum after binning and denoising with the polynomial baseline overlay; the lower plot shows the final baseline-subtracted and normalised spectrum. Use **Previous/Next** to navigate between files, and **Save** to export the current result. **Switch to Batch Process** transitions to the Spectrum Batch Processing module.

**Fig 18:**
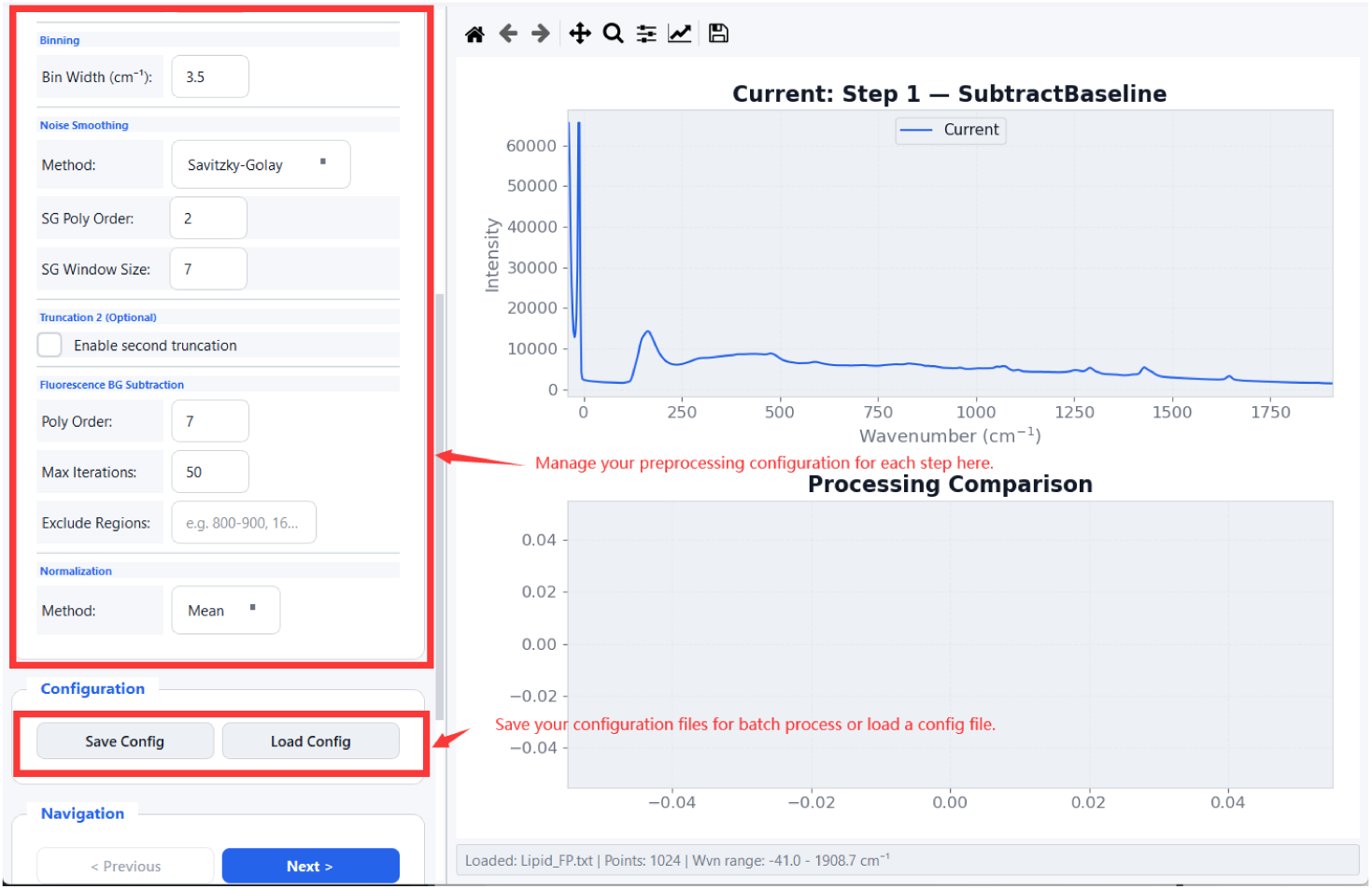
Processing Parameters panel: pipeline steps are grouped with labelled parameters for Truncation, Binning, FBS, Normalization, and Noise Smoothing. Configuration can be saved and reloaded.

**Fig 19:**
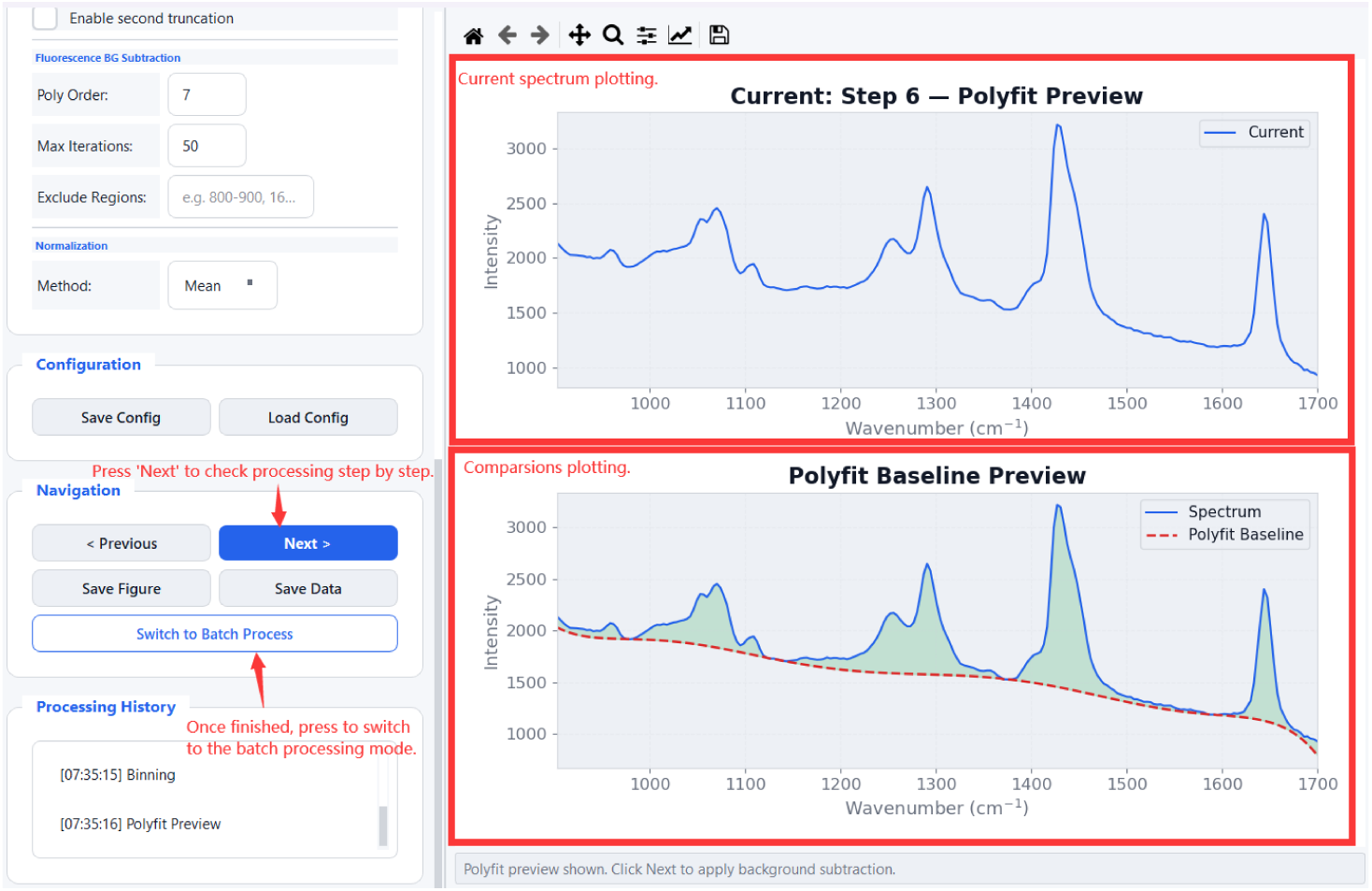
Spectrum Processing result view: dual plot showing the denoised spectrum with fitted polynomial baseline (top) and the final processed spectrum (bottom). Navigation and save controls are shown at right.

### A.5 Spectrum Batch Processing

Spectrum Batch Processing applies the same pipeline to a collection of spectra without manual intervention. The workflow mirrors the single-spectrum mode but operates on multiple files simultaneously.

#### Loading configuration

In the Processing Parameters panel (Figure 20), click **Load** to import a previously saved JSON configuration file. The Processing Log panel (right) confirms the loaded configuration path. All parameter fields remain editable after loading.

**Fig 20:**
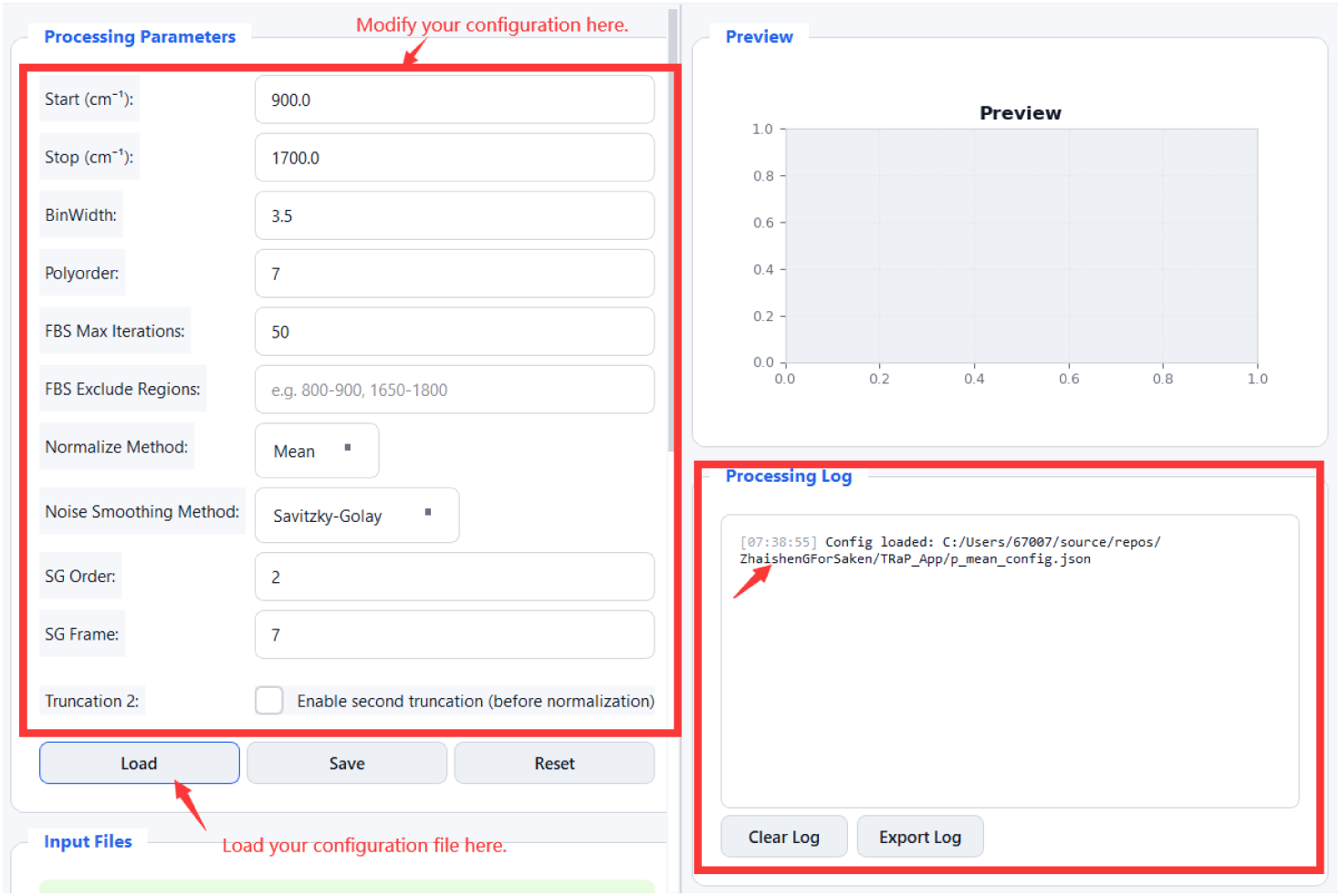
Spectrum Batch Processing: loading a configuration file. The Processing Log panel confirms the loaded config path. Parameters shown include Start/Stop, BinWidth, Polyorder, FBS Max Iterations, Normalize Method, and Noise Smoothing settings.

#### Input Files and Actions

In the Input Files panel (Figure 21), click **Select Data Files (Raw Spectrum)** to choose multiple files at once (the list shows all selected filenames with a count). Then provide the WL correction file, calibration `.mat` file, output folder (defaults to `./Processed` relative to the data location), and an optional subfolder name. Click **Preview First File** to verify parameters on a single spectrum before committing, then click **Start Batch Process** to process all files.

**Fig 21:**
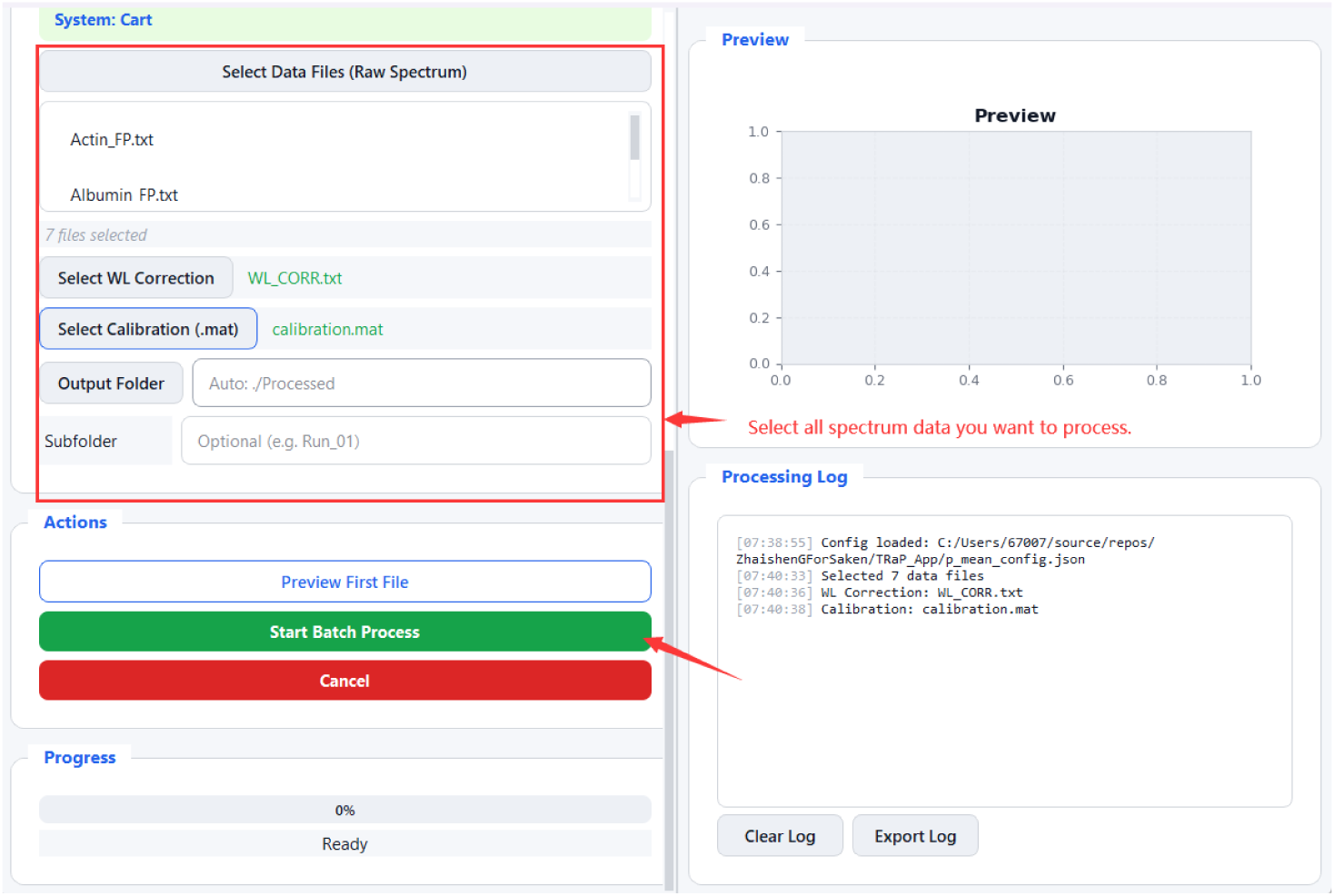
Spectrum Batch Processing Input Files panel: 7 spectra selected, WL correction and calibration files assigned, output folder configured. *Preview First File* validates settings; *Start Batch Process* processes all files.

#### Completion

Upon completion, a summary dialog reports the number of successfully processed and failed files (Figure 22). The Processing Log lists each file’s output filename, which encodes the key processing parameters. The log can be exported for record-keeping using **Export Log.**

**Fig 22:**
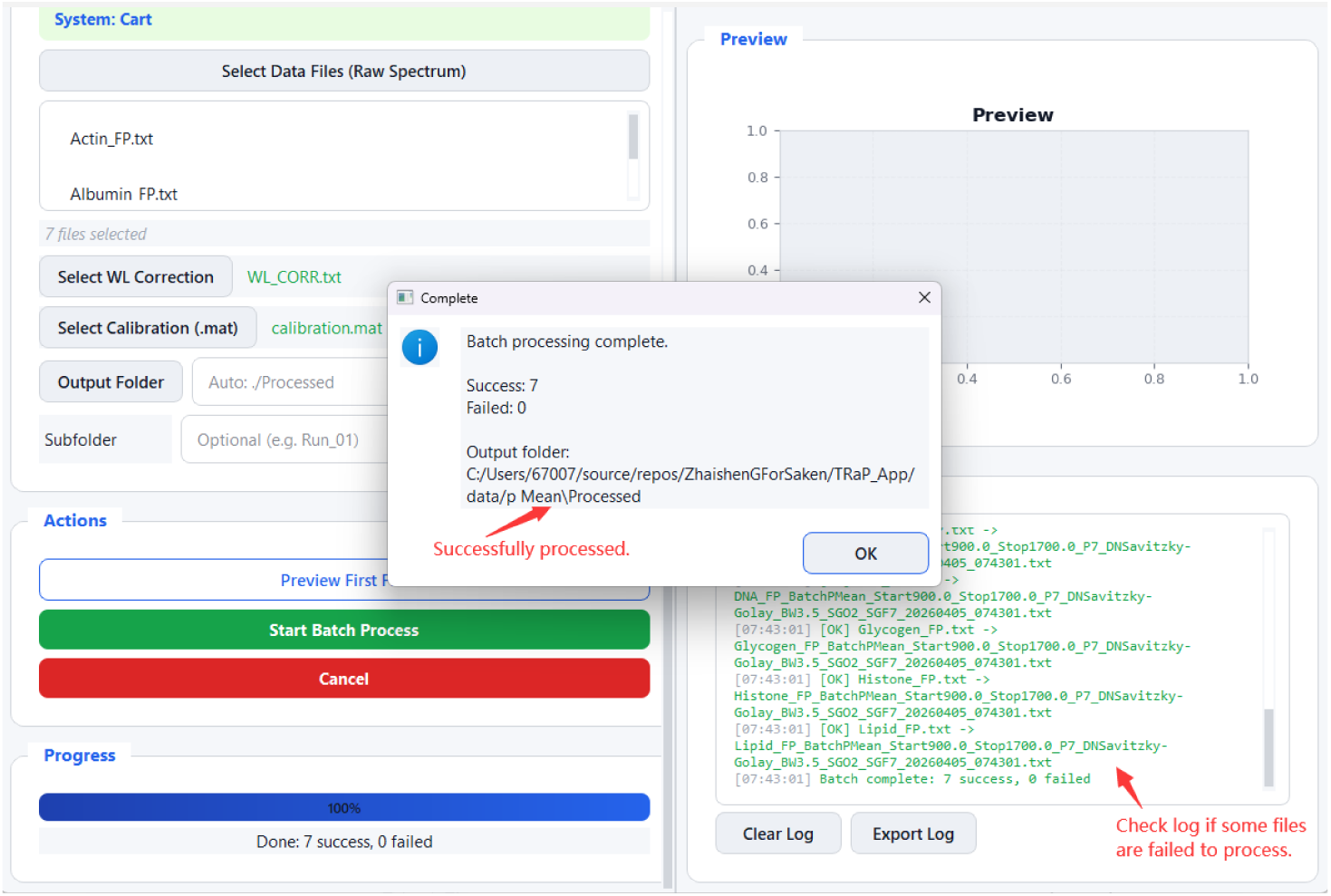
Spectrum Batch Processing completion: summary dialog reports 7 successes and 0 failures. The Processing Log lists each output filename with embedded processing parameters and timestamp. Failed files, if any, are identified in the log for inspection.

